# Glyphosate used to control invasive *Phragmites australis* in standing water poses little risk to aquatic biota

**DOI:** 10.1101/2020.06.19.162222

**Authors:** C.D. Robichaud, R.C. Rooney

## Abstract

When an invasive wetland grass degrades a Ramsar wetland and Important Bird Area, decisive management action is called for. To limit the extent and spread of European *Phragmites australis*, the Ontario government began the first, large-scale application of glyphosate (Roundup Custom®) over standing water to control an invasive species in Canadian history. Between 2016 and 2018, over 1000 ha of marsh were treated. To assess the risk this herbicide presented to aquatic biota, we measured the concentration of glyphosate, its primary breakdown product aminomethylphosphonic acid (AMPA), and the alcohol ethoxylate-based adjuvant Aquasurf® in water and sediments in areas of the highest exposure risk and up to 150 m into adjacent bays. We never detected glyphosate or AMPA at concentrations exceeding thresholds of toxicological concern. The maximum observed concentration of glyphosate in water was 0.320 ppm, occurring within 24 hr of application. The maximum glyphosate concentration in sediment was 0.250 ppm, occurring within 30 days of application. AMPA was detectable in water and sediment, indicating microbial breakdown of glyphosate in the marsh, but at low concentrations (max_water_ = 0.025 ppm, max_sed_ = 0.012 ppm). The maximum distance from the point of application at which glyphosate was detected in the water was 100 m, vs. 0 m for AMPA. Concentrations in water returned to pre-treatment levels (<DL) within 20-30 days of application. In sediment, glyphosate residue persisted above detection limits (>0.005 ppm) for over one year but less than two years. Concentrations of alcohol ethoxylates were variable in space and time, following a pattern that could not be attributed to Aquasurf® application. The direct, over-water application of Roundup Custom® with Aquasurf® to control invasive *P. australis* does not pose a toxicological risk to aquatic biota.

**Highlights:**

1. Glyphosate-based herbicide was applied directly to >1000 ha of marsh to control invasive *P. australis*
2. Glyphosate and AMPA did not reach levels of toxicological concern for aquatic biota
3. Aquasurf® exceedances were observed but could not be attributed to *P. australis* control activity
4. Glyphosate, AMPA, and Aquasurf® dispersed no more than 100 m from the point of application
5. Glyphosate, AMPA and Aquasurf® in water returned to baseline levels within 30 days of application
6. Glyphosate, AMPA and Aquasurf® in sediment returned to baseline levels within 2 years

## Introduction

Glyphosate (*N-*[phosphonomethyl]glycine), a broad-spectrum, post-emergence herbicide (Solomon and Thompson, 2003), is the most heavily used pesticide in North America (Benbrook, 2016). In Canada, over 2900 tonnes was applied in Ontario in 2013 alone, mainly on corn and soybeans (Farm & Food Care Ontario, 2015). Due to its extensive use, glyphosate and its primary breakdown product aminomethylphosphonic acid (AMPA) are often detected in ground and surface water (Struger et al., 2008; Van Stempvoort et al., 2014). It can reach aquatic ecosystems, such as wetlands, through a combination of wet and dry aerial deposition, accidental overspray, and run-off from agricultural landscapes (Glozier et al., 2012; Montiel-León et al., 2019; Struger et al., 2015).

Once it reaches wetlands, glyphosate (and subsequently AMPA) may present a risk to biota (Annett et al., 2014), though environmental concentrations are typically below thresholds of toxicological concern (Byer et al., 2008; Struger et al., 2008). While the risk of exceeding toxicological thresholds for glyphosate and AMPA in wetland ecosystems may be low when transported by drift, run-off or accidental overspray, when glyphosate is applied directly to the wetland to control invasive plants the risk to aquatic biota is likely much greater, though this has not been well studied (Breckels and Kilgour, 2018).

Invasive wetland plants (e.g. *Phragmites australis, Phalaris arundinacea, Typha* x *glauca, Arundo donax*) tend to share traits that make them difficult to eradicate, including rhizomatous growth (Zedler and Kercher, 2004). Physical removal methods (i.e. cutting, rolling, burning, pulling) are not effective on their own because they do not kill belowground rhizomes, allowing the invasive plants to persist (e.g., Derr, 2008). In comparison, glyphosate treatment on its own or in combination with mechanical control can be highly effective because it affects both the above and belowground portions of the plant (e.g., OMNRF, 2011).

While many invasive wetland plant control projects employ glyphosate-based herbicides (e.g., Kettenring and Adams, 2011; Martin and Blossey, 2013), the application of glyphosate to control invasive wetland plants is limited in Canada by the lack of a registered product to use over standing water (Annett et al., 2014; CCME, 2012). This has substantially hindered efforts to control the aggressive perennial grass *Phragmites australis* ssp. *australis* (European Common Reed ((Cav.) Trin. Ex Steud); henceforth *P. australis*), limiting herbicide-based control to areas where there is no standing water (e.g., Crowe et al., 2011). *Phragmites australis* has a wide range of environmental tolerances and has become widespread in Canada (Catling and Mitrow, 2011). Because it can grow in anything from moist soils to over 1.5 m of standing water (Hudon et al., 2005), restricting herbicide application to areas where the water table is below ground has left remnants from which *P. australis* rapidly recolonizes treated areas (e.g., Lombard et al., 2012; Quirion et al., 2017).

*Phragmites australis* creates dense monocultures that demonstrably reduce habitat quality for at-risk amphibians (Greenberg and Green, 2013), turtles (Markle and Chow-Fraser, 2018), and birds (Robichaud and Rooney, 2017). It is listed as a threat to 25% of Ontario’s species at risk, particularly plant species at risk (Bickerton, 2015). Because of the direct threat to species of conservation concern, and consequent impairment of ecological integrity, the Ontario Ministry of Natural Resources and Forestry obtained approval from Health Canada’s Pest Management Regulatory Agency for an Emergency Registration as a pilot project to apply glyphosate in areas with standing water in Rondeau Provincial Park and the Long Point region, on the North Shore of Lake Erie (MNRF 2018). This project was the first large-scale (over 1165 ha treated) use of glyphosate to control invasive *P. australis* in standing water in Canada. Due to the unprecedented scale and the Emergency Registration required to enable the control project, our research team designed and implemented extensive monitoring to evaluate the fate and potential effects of the herbicide (MNRF, 2018; Yuckin and Rooney, 2019; Robichaud and Rooney, *in prep*; Polowyk and Rooney, *in prep*).

As part of this monitoring effort, we sought to determine if direct, over water application of glyphosate-based herbicide led to concentrations of glyphosate, AMPA, or alcohol ethoxylates from the associated surfactant (Aquasurf®) in sediment or surface water that could pose a risk to aquatic biota. Second, we wanted to assess dispersal of glyphosate, AMPA, and Aquasurf® from the point of application. Third, we wanted to evaluate the persistence of glyphosate, AMPA and Aquasurf® in the water and sediment following application. Our results fill a key knowledge gap about the potential risks of direct application of glyphosate over standing water to control an invasive wetland plant, a practice not currently legal in Canada outside of the Emergency Registration process.

## Methods

### Study area

The pilot *P. australis* control project took place in three hemi- and emergent marsh complexes on the north shore of Lake Erie: Rondeau Provincial Park (Fig. 1A), Long Point (Fig. 1A) and Turkey Point (Fig. 1B). These marsh complexes are ecologically significant in the surrounding landscape and are home to many at-risk and provincially rare species (Ball et al., 2003). Long Point is a UNESCO Biosphere Reserve and a Ramsar wetland of international importance, and both Rondeau Provincial Park and Long Point are designated Important Bird Areas that provide essential habitat for migratory and marsh-nesting bird populations. *Phragmites australis* rapidly expanded in these marsh complexes during the 1990s (Wilcox et al., 2003) and is predicted to expand at 14 to 37% per year into 2022 without active intervention (Jung et al., 2017).

**Figure 1.**
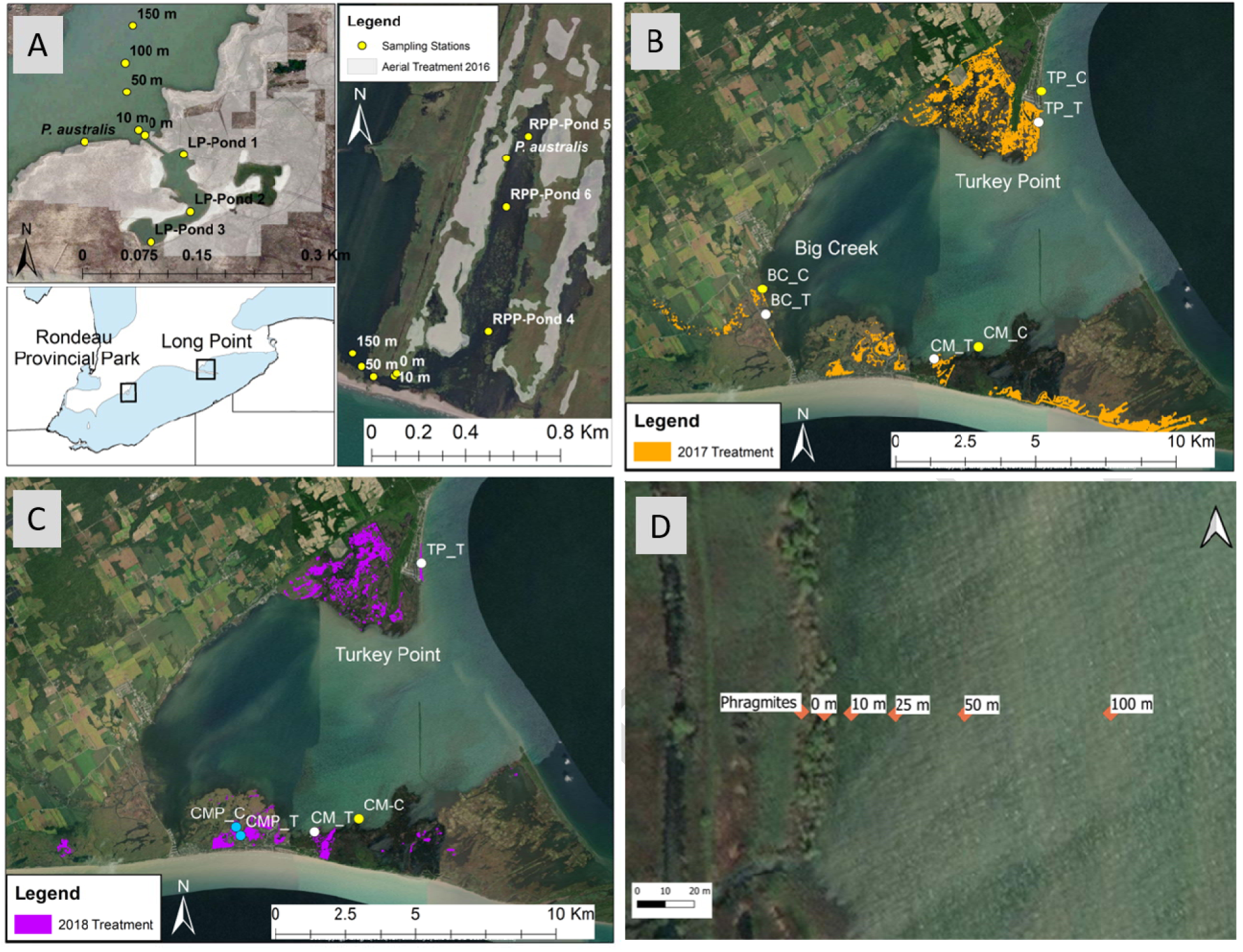
Control (C) and treatment (T) sampling stations in Long Point (LP) and Rondeau Provincial Park (RPP) in Ontario, CA in 2016 (A), and in the Long Point region in 2017 (B), and 2018 (C). To evaluate glyphosate, AMPA, and Aquasurf® concentrations we established sites in areas of maximum exposure (ponds, *P. australis*, 0 m stations). To assess dispersal from the point of application we established transects stretching from points of maximum exposure into the adjoining bay (example in panel D). Sampling occurred on three occasions at each station: a pre-treatment timepoint, within 24 hours of herbicide application, and >20 days post-application. Areas of dense *P. australis* (illustrated by coloureds polygon: white in 2016, orange in 2017, purple in 2018) were treated with a glyphosate-based herbicide: approximately 100 ha in Rondeau PP and 355 ha in Long Point in 2016, 329 ha in 2017, and 99.89 ha in 2018. Image from Google Earth. Google Earth V 7.3.2.5776. 2019. TerraMetrics 2019.

### Treatment plan

Details regarding the annual treatment plans are published in annual reports (MNRF 2016, 2017, 2018). In brief, between 2016 and 2018, in partnership with the Nature Conservancy of Canada, the Ontario Ministry of Environment, Conservation and Parks, and the Canadian Wildlife Service, the Ontario Ministry of Natural Resources and Forestry obtained an Emergency Registration (#32356) under the Pest Control Products Act from Health Canada’s Pest Management Regulation Authority and a provincial Permit to Perform an Aquatic Extermination, which allowed them to apply a glyphosate-based herbicide (Roundup® Custom For Aquatic & Terrestrial Use Liquid Herbicide, Registration Number 32356 Pest Control Products Act) combined with a non-ionic alcohol ethoxylate surfactant (Aquasurf®, Registration Number 32152) to control invasive *P. australis* in standing water within the study area.

In 2016, contractors working for the aforementioned partners conducted herbicide treatment primarily via helicopter in Rondeau Provincial Park and the Long Point region. One-hundred hectares of marsh in Rondeau and approximately 355 ha in Long Point were treated with 4210 g acid equivalent glyphosate/ha as an isopropylamine salt (Roundup Custom®), combined with Aquasurf® non-ionic surfactant at 0.5 L/ha. This herbicide mix was applied at 8.77 L/ha, with a total spray mix of 70 L/ha, by a Eurocopter A-Star that was equipped with GPS guidance and auto-booms with Accu-flo nozzles. The aircraft was calibrated to consistently deliver droplets in the ASABE coarse to very-coarse range (ASABE S572.1) with a maximum spraying speed of 60 km/hour delivered from a maximum of three metres above the plants. Treatment took place between the 6th and 23rd of September when wetland flora had begun to senesce and faunal breeding seasons were complete to limit the impact of management on wetland species, following best management practices (OMNRF 2011).

In 2017, 329 ha of marshland in the Long Point region were treated by ground application of the same blend of Roundup Custom® and Aquasurf® surfactant, but at a reduced loading rate of 1200-3600 g acid equivalent glyphosate/ha with a 0.5-1.5% solution of Aquasurf® added at a percentage-by-volume basis to achieve desired concentrations. Ground treatment was completed using a combination of boat and an amphibious Marsh Master™. As only 17 ha were treated in Rondeau Provincial Park, our monitoring focused on Long Point region (Fig. 1B).

In 2018, approximately 100 ha of herbicide application took place in the Long Point region (Figure 1C), again delivered primarily via ground application. This treatment was a combination of applying herbicide to un-treated *P. australis* as well as follow-up in previously treated locations. Though follow-up treatment involves applying the same herbicide mixture as is used in the original treatment, the loading rate is lower on an area basis because herbicide is only applied to patches of *P. australis* that either resisted control in previous years or managed to recolonize.

### 2016 Field methods

To evaluate the maximum exposure, dispersal, and persistence of glyphosate, AMPA, and Aquasurf® surfactant, we established nine sampling stations in both Long Point and Rondeau Provincial Park (Figure 1A). At each project site, three stations were placed within a semi-isolated pond surrounded by *P. australis* that was to be treated by aerial application of herbicide in 2016, and one within a patch of *P. australis* that was also to be herbicide-treated. To evaluate the spatial extent and movement of analytes from the point of application, we established transects stretching into Long Point and Rondeau Bays, each consisting of five stations. The transects began at the mouth of the semi-isolated ponds (0m), then had sampling stations located 10 m, 50 m, 100 m, and 150 m into their respective bay (Figure 1A). At each sampling station we collected a depth-integrated water column sample using a plexiglass tube and a sample of the top 10 cm of sediment with a Wildco stainless-steel Petite Ponar Grab sampler (0.023 m^2^). The water column sampler and Ponar were rinsed three times between stations to prevent cross contamination.

All stations were sampled three times: 1) at a pre-treatment time point before herbicide application that year, 2) within 24 hours of herbicide application to capture maximum exposure levels, and 3) >20-days after application. For exact dates see Appendix A.

### 2017 Field methods

Ground application took place primarily along the margin of Long Point Bay, therefore our 2017 sampling design comprised transects extending into the Long Point Bay. To clarify the effects of herbicide exposure we paired ‘control’ and ‘treatment’ transects, with control transects established as a reference such that they were within 2 km but greater than 1 km away from the treatment transect. We thus established six transects in three management units: Big Creek National Wildlife Area, Turkey Point Provincial Park, and Crown Marsh (Figure 1B). Transects consisted of six stations – the first within *P. australis* growing on the edge of the Bay, then 0 m, 10, 25, 50, and 100 m from the shoreline (Figure 1D). As in 2016, sampling occurred pre-treatment, within 24 hour of herbicide application, and >20-day time points (SM1). Note that because ground-based application is slower than aerial application, treatment in 2017 continued for a period of a week. The water and sediment field sampling followed 2016 methods.

To evaluate longer-term persistence of glyphosate, AMPA, and Aquasurf® surfactant we re-sampled a subset of the 2016 stations from Rondeau Provincial Park and Long Point. In August 2017, before treatment began anywhere else in the region, we collected water and sediment at the *P. australis* station, the central pond station, and the 0 m, 10 m, and 50 m transect stations.

### 2018 Field methods

A combination of new treatment and follow-up treatment occurred in 2018. New treatment took place in Turkey Point and Crown Marsh, while follow-up treatment took place in a different region of Crown Marsh. To monitor the newly treated *P. australis* sites we established three sites in one control and one treatment pond adjacent to new treatment in Crown Marsh (n = 6) and one transect in Turkey Point (Fig. 1C). The Turkey Point treatment was in the location of the 2017 control transect, so we re-sampled the stations from within the *P. australis* extending to 50 m along this transect in 2018. There was no suitable location for a paired control transect, so sampling consisted of only the treatment transect in 2018.

The 2018, follow-up treatment in Crown Marsh occurred in the location of our 2017 treatment transect - therefore, we re-sampled the treatment and control transects established in 2017, stretching from in *P. australis* out 100 m into the Long Point Bay (Fig. 1D). Finally, in September 2018, to continue our assessment of the long-term persistence of glyphosate, AMPA and Aquasurf® surfactant residue we re-sampled three stations in the Crown Marsh pond that was treated in 2016.

### Lab analyses and ecotoxicology thresholds

Water and sediment samples were submitted to the University of Guelph Agriculture and Food Laboratory for analysis. Glyphosate, AMPA and alcohol ethoxylate concentrations in water and sediment were determined using their LC-MS/MS chromatography method. Laboratory samples are homogenized so that a 5–100 g test portion taken for analysis is representative of the entire sample. For water samples, the sample is evaporated and reconstituted into methanol, whereas sediment samples are extracted into a 10% acidified water and methanol solution and separated from co-extractives using solid phase extraction. The solution is then passed through an analytical column. For glyphosate and AMPA, the LC-MS/MS system employs a cation guard column (Micro-Guard Cation-H cartridge 30 × 4.6 mm) for chromatographic separation, with 0.1% formic acid in nanopure grade water as mobile phase A and acetonitrile as mobile phase B. The mass spectrometer employed is a SCIEX 4000 with a negative polarization. For alcohol ethoxylates, in 2016 and 2017 the analytical column used for chromatographic separation was a ZIC HILIC 100 x 4.6 mm, but this was updated to a ZIC HILIC 50 x 2.1 mm column in 2018 for improved peak resolution. In both cases, mobile phase A comprised 0.4% formic acid in 4 mM ammonium formate and mobile phase B was 5 mM ammonium acetate in methanol. The mass spectrometer employed in alcohol ethoxylate analyses was an AB Sciex triple Quad 5500 with positive polarization.

Of note, the concentration of alcohol ethoxylates reported includes the sum of homologues C_12_EO_6,_ C_12_EO_7_, C_12_EO_8_, C_12_EO_9_, C_13_EO_6_, C_13_EO_7_, C_13_EO_8_, and C_13_EO_9,_ which analysis of the Aquasurf® surfactant indicated were the homologues best representative of the hundreds of homologues present in the adjuvant. Thus, it provides an estimate of the total alcohol ethoxylates in the system but does not include all homologues that are present in Aquasurf®. The homologue C_12_EO_6_ was the most sensitive ion to detection and was used as the fundamental quantitative peak when determining alcohol ethoxylate concentrations (Agriculture and Food Laboratory, University of Guelph pers comm.).

**Table 1.** Limits of detection and limits of quantification for glyphosate, aminomethylphosphonic acid (AMPA) and alcohol ethoxylate homologues (AEH) most representative of Aquasurf® surfactant in this study. Samples with values between the limit of detection and limit of quantification have estimated values reported, as they represent certainty around the presence of the chemical residue but uncertainty about the exact concentration. Analyses took place at the Agriculture and Food Laboratory at the University of Guelph.

| Analyte | Matrix | Limit of detection (ppm) | Limit of quantification (ppm) |
| --- | --- | --- | --- |
| **Glyphosate** | Water | 0.001 | 0.008 |
|  | Sediment | 0.005 | 0.020 |
| **AMPA** | Water | 0.002 | 0.008 |
|  | Sediment | 0.005 | 0.020 |
| **Aquasurf® surfactant** | Water | 0.030 | 0.080 |
|  | Sediment | 0.300 | 0.900 |

To assess the relationship between the herbicide and turbidity, we collected total suspended solids (TSS) (mg/L) from each station every year, excluding the two 2018 *P. australis* sites where the water was too shallow to obtain a clean sample (n = 46). Total suspended solids were measured by weighing the residue from a 500 mL sample of water captured by a gf/f Whatman filter paper (0.7 micron pore size), following ESS Method 340.2 (EPA, 2003). A subset of stations over the three years also had sediment analyzed for iron (Fe) (mg/kg dry). Samples were extracted using a 0.005M DPTA solution, and the filtrate was analysed by Varian ICP-OES (Liang and Karamanos, 1993) by the Agriculture and Food Laboratory in Guelph, ON. Iron was measured at each station in 2016, at the 0 m Crown Marsh and Turkey Point stations in 2017, and the 0 m Crown Marsh station in 2018 (n = 21).

### Data analysis

To evaluate the maximum exposure risk resulting from *P. australis* control efforts, concentrations of glyphosate, AMPA, and Aquasurf® surfactant measured in water and sediment samples were compared to available ecotoxicology thresholds. The Canadian Council of Ministers of the Environment (CCME) has set maximum exposure thresholds for short-term (27.000 ppm) and long-term exposure (0.800 ppm) to glyphosate in freshwater (CCME, 2012), which we also apply to AMPA. For alcohol ethoxylates we employed the Human and Environmental Risk Assessment (HERA) guidelines, which provided Predicted No Effect Concentrations (PNECs) for water and, based on partitioning coefficients, extrapolates these to sediment (HERA, 2009). We selected the homologue C_12_EO_6_ for our toxicity threshold, as this was the most sensitive AE to detection in our samples. The HERA report suggests a PNEC_water_ for the AE homologue (C_12_EO_6_) of 0.129 ppm. The 2013 draft Canadian Federal Environmental Quality Guideline for AEs by homologue for C_12_EO_6_ has a water quality guideline value of 0.193 ppm. The PNEC_soil_ for (C_12_EO_6_) is 25.100 ppm and is calculated based on chronic *Daphnia magna* PNEC_water_ values, using an equilibrium partitioning method (HERA, 2009). In the case of glyphosate in sediment, there is no established threshold for safe exposure in Canada.

Consequently, we applied the HERA PNEC_soil_ calculations for alcohol ethoxylate homologues to glyphosate. Using the average fraction of organic carbon in soil from our 2017 study sites and the long-term exposure to glyphosate guidelines for protection of aquatic life (0.800 ppm), we estimated a threshold of 206.720 ppm (HERA, 2009; equations TGD 72, 24, 23).

The dispersal distance for each analyte within 24 h of application was identified as the maximum distance beyond the point of application where the analyte was detected above pre-treatment levels. To identify the persistence of elevated analyte concentrations, the concentration of analytes following application was evaluated and compared to pre-treatment concentrations at all sampled stations at >20 days post-application and at some stations for up to two years after application.

Because glyphosate is known to adsorb rapidly to suspended sediment (Franz et al., 1997), the concentrations of glyphosate and AMPA in water within 24 hours of application were compared to total suspended solids (mg/L) (TSS) (n = 46). Similarly, we assessed how total suspended solids (TSS) and Fe (mg/kg) influenced concentrations of glyphosate and AMPA detected in sediment samples within 24 hours of application (n = 21). Aquasurf® was not analyzed because there were too few occurrences in the dataset. To evaluate these relationships, we used linear mixed models with station, or distance from the site of application, as a random. Linear mixed models were conducted using the *lme4* package (Bates et al., 2015). Total suspended solids and iron were both log_10_ transformed before analysis. We determined the conditional (r^2^_c_) and marginal (r^2^_m_) coefficient of variation using the approach specified by Nakagawa and Schielzeth (2013) with the *MuMIn* package (Barton, 2020). The conditional r^2^ reports the variation explained by the entire model, while the marginal r^2^ reports the variation attributed to the fixed effects. We compared our model with a null model (intercept, with distance as a random factor) using a likelihood test ratio (Bolker et al., 2009; Pinheiro and Bates, 2000) via the *car* package (Fox and Weisberg, 2019). And finally, we conducted model comparison using AICcmodavg (Mazerolle, 2019). All analyses were performed in R v 3.6.2 (R Core Team, 2016). All figures were made using ggplot2 (Wickham, 2016). In all figures and analyses, concentrations that were below the limit of detection (BDL) were treated as zeros.

## Results

### Maximum exposure risk

There were no concentrations of glyphosate or AMPA that approached the thresholds set for the protection of aquatic biota over the three years. Stations considered to be at maximum exposure risk (i.e. ponds, the *P. australis* sites, and the 0 m transect station) had concentrations of glyphosate and AMPA in water that peaked immediately following herbicide application, (Fig. 2B, Appendix B), whereas glyphosate concentrations in sediment were highest at the >20 day time point (Fig. 2F). The concentrations of AMPA in sediment were all below the limit of quantification (Appendix B). The maximum concentrations detected were 0.320 ppm glyphosate and 0.025 ppm AMPA in water (Appendix C) and 0.250 ppm glyphosate and 0.012 ppm AMPA in sediment (Appendix D).

**Figure 2.**
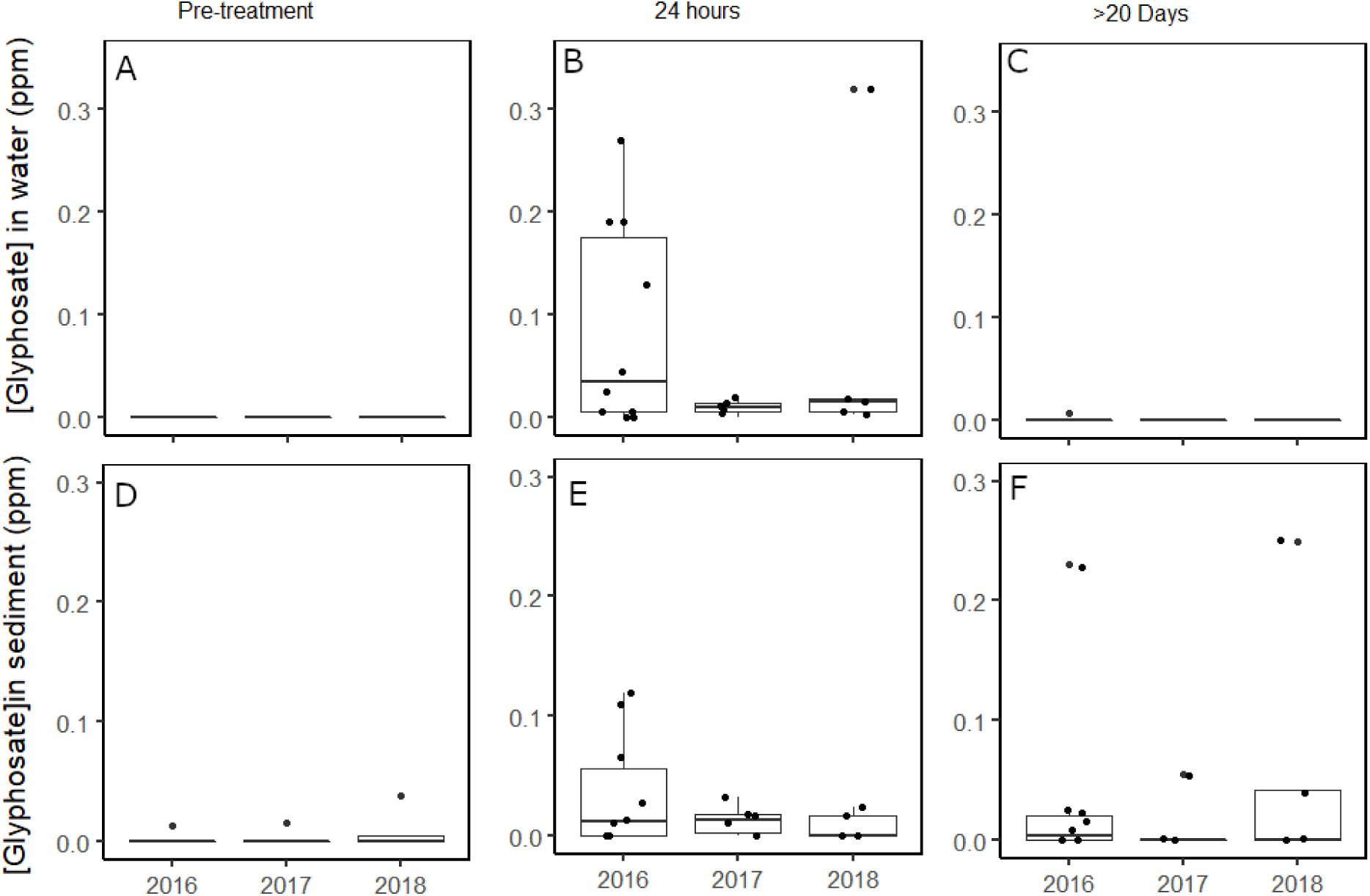
Concentrations of glyphosate in water (A-C) and sediment (D-F) at sites with maximum exposure risk (i.e. *P. australis*, ponds, and 0 m transect sites) from treatment transects in Rondeau Provincial Park, Long Point, Big Creek National Wildlife Area, and Turkey Point, Ontario, CA (2016: n = 10, 2017: n = 6, 2018: n = 5). Boxplots represent the median, the 25th (lower hinge) and 95th (upper hinge) percentiles and 1.5*IQR (upper/lower notch).

Detections of alcohol ethoxylates from the Aquasurf® surfactant were infrequent and did not conform spatially or temporally to a pattern that could be attributed to the herbicide treatment. Within 24 hours of application in 2016 and 2017, alcohol ethoxylates were only detected at stations situated in areas where herbicide was applied directly, and always at concentrations below thresholds of toxicological risk (Appendix C & D). In 2018, there were few detections at the herbicide transect within 24 hours, with the highest concentration at a *P. australis* site (0.270 ppm) (Appendix E & F). There were detections of Aquasurf® in sediment within 24 hours of treatment at the *P. australis* sites (Appendix E & G), and one of these detections was at a control transect where no treatment had occurred.

### Dispersal from point of application

Within 24 hours of herbicide application, concentrations of glyphosate, AMPA and alcohol ethoxylates were highest at the stations closest to and within the treated *P. australis*. Of the three chemicals of concern, we observed glyphosate dispersing the furthest within 24 hours of treatment to a maximum distance of 100 m in water and 50 m in sediment in trace concentrations (Appendix F & G). In comparison, AMPA was not detected, in either water or sediment, past the 0 m transect station (Appendix F, G & H). AMPA always co-occurred with glyphosate, with the exception of one sample from a highly flushed 0 m transect station, sampled >20 days after treatment in 2018, where AMPA exceeded the detection limit (0.005 ppm), but not the limit of quantification (0.02 ppm).

As described in the section on maximum exposure risk, the detection of alcohol ethoxylates in water and sediment did not conform to any anticipated pattern. For example, in 2017 alcohol ethoxylates were only detected in one sediment sample, but this sample was collected prior to any herbicide treatment (pre-treatment). Also, in 2017, alcohol ethoxylates were detected in a single water sample that was collected approximately a month after herbicide application at a station 100 m from the nearest point of application. In 2018, at the >20 days timepoint, alcohol ethoxylates were detected in water from every station along the Crown Marsh treatment transect, except for the *P. australis* station, despite no detections in water within 24 hrs of application. More, alcohol ethoxylates were simultaneously detected in water from nearly every station along the Crown Marsh control transect, where no herbicide was applied (Appendix I).

### Persistence in water and sediment

Prior to treatment, water had no detectable levels of glyphosate (Fig. 2A) or AMPA (Appendix B). Alcohol ethoxylates were also below detection levels in all water samples collected pre-treatment, except for one 50 m transect station in 2018. Concentrations of glyphosate (Fig. 3C) and AMPA (Appendix B) in water returned to pre-treatment levels by the >20 days after treatment timepoint, except one *P. australis* site in Rondeau Provincial Park (Fig. 2C, Fig. 3C). Upon resampling in 2017 and 2018 no glyphosate or AMPA was detected in water samples (Fig. 3D, Appendix H).

**Figure 3.**
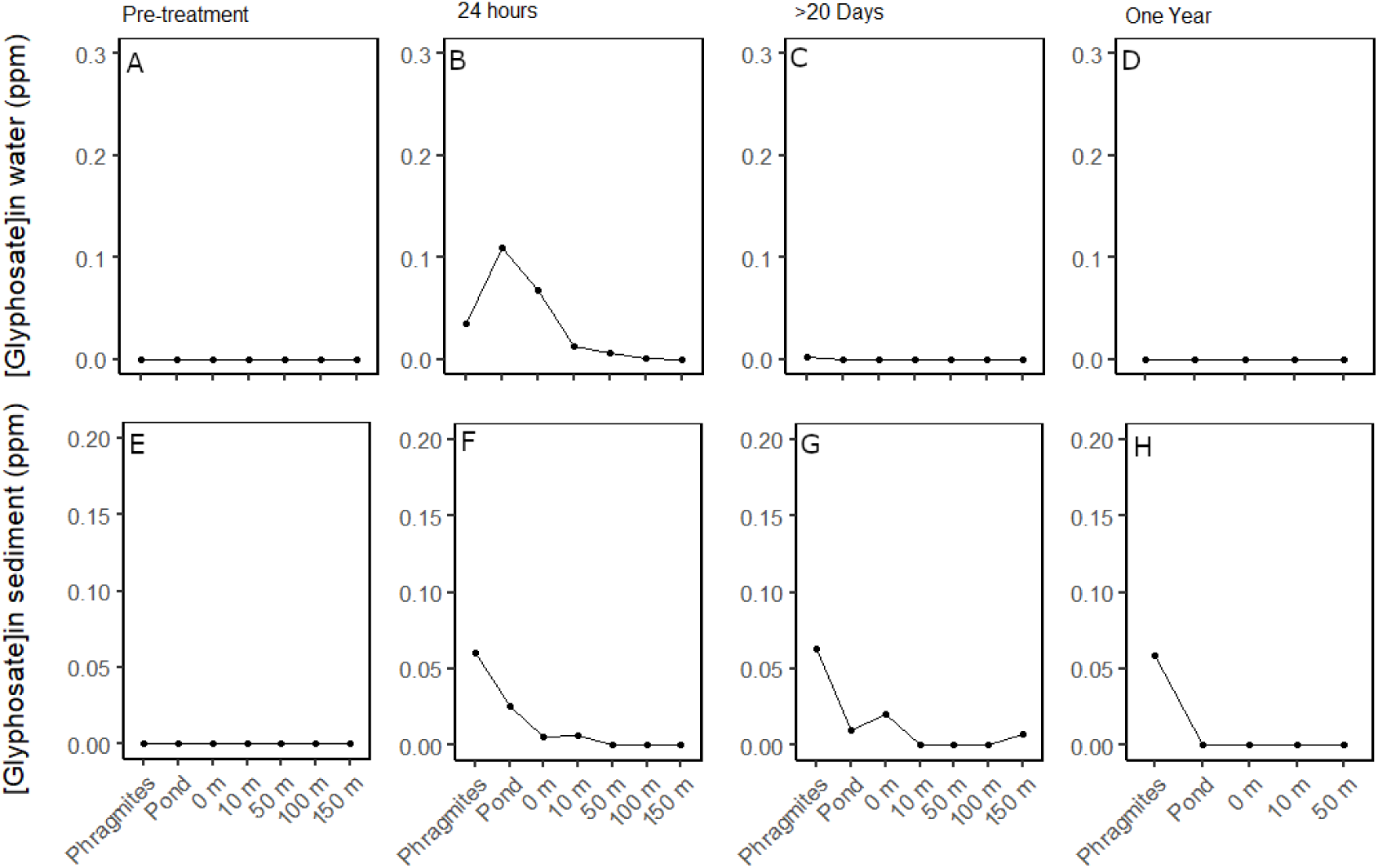
Concentrations of glyphosate in water and sediment samples collected from Rondeau Provincial Park and Long Point, Ontario, CA. Samples were collected before aerial-application of glyphosate-based herbicide, within 24 hours, >20-days after, and one-year after application. Sites represent areas of maximum exposure to herbicide (*P. australis*, ponds, 0 m) and distance (m) out into the adjoining bay. The average concentration at each point is represented here (pond: n = 6; Phragmites, 0, 10, 50, 100 and 150 m: n = 2). Data was collected in 2016 and 2017 and is presented in Appendices C, D, F.

As mentioned earlier, alcohol ethoxylates were present at the >20-day timepoint, but whether they are a result of the herbicide application is difficult to determine. In 2016, one detection of alcohol ethoxylates in water occurred at the *P. australis* site at the >20-day timepoint in Rondeau (Appendix E). In 2017, trace amounts of alcohol ethoxylates (0.004 ppm) were detected in water collected >20-days after herbicide application at the 100 m treatment transect in Crown Marsh (Appendix D). And in 2018, alcohol ethoxylates were detected in water from five of the six treatment transect stations and four of the six control transect stations >20-days after treatment in Crown Marsh, but at none of the Turkey Point transect stations on the same date (Appendix I). When the 2016 station was resampled in 2017 and 2018, no alcohol ethoxylates were detected in the water (Appendix J).

Glyphosate and AMPA were more persistent in sediment than in water. The highest observed concentrations of glyphosate in sediment occurred at the >20-day post application timepoint (Fig. 2F, Appendix D). Trace amounts of glyphosate were detected as far as 150 m from the point of application at the >20-day timepoint in 2016, though this value was below the limit of quantification. In August 2018, when we resampled the 2017 treatment transect stations at Crown Marsh, we detected low levels of glyphosate in sediment at the *P. australis* and 0 m transect stations at the pre-treatment time point, indicating residue had persisted from the previous treatment (Fig. 4). Similarly, in August 2017, when we resampled a subset of stations, we found detectable glyphosate and AMPA at one station in Crown Marsh where concentrations had been highest in 2016 (Fig. 3H, Appendix H). When we surveyed the sediment at these locations again in November 2017, glyphosate persisted in sediment while AMPA was below detection limits. By September 2018, two years after treatment, glyphosate in the sediment was also below the detection limit (Appendix J). Thus, we conclude that glyphosate can persist in sediment at detectable levels for at least one-year post-treatment. Alcohol ethoxylates, in contrast, were rarely detected in the sediment. They were only observed in sediment at one station, the 25 m transect station in Turkey Point, during the >20 days timepoint in 2018.

**Figure 4.**
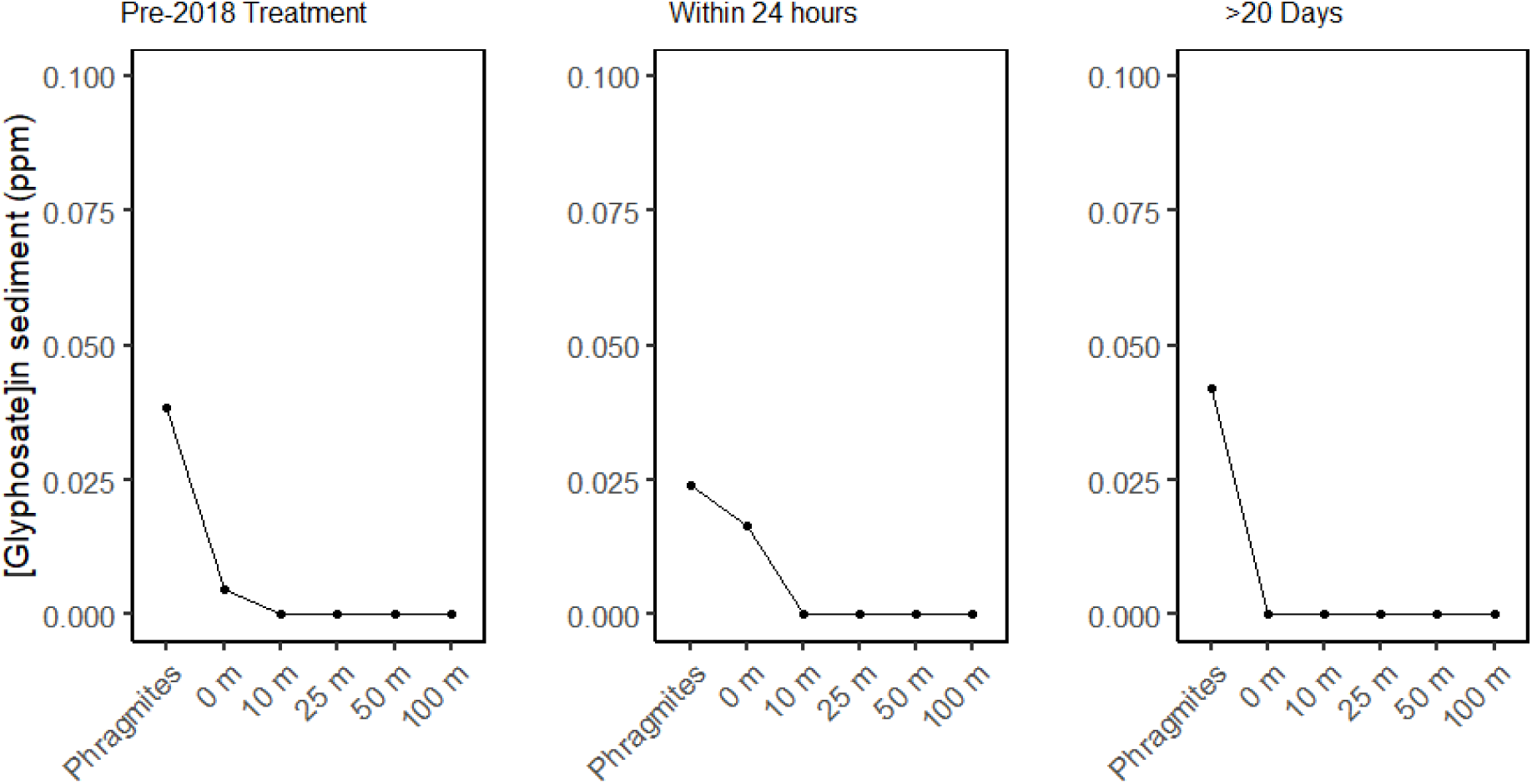
Glyphosate residue remained in sediment one year after treatment in Crown Marsh, Long Point (Ontario, CA). The transect was treated with a glyphosate-based herbicide in 2017 and glyphosate was present at the pre-treatment timepoint in 2018. These concentrations are below thresholds of concern for aquatic biota.

Of note, in 2016, there was a detection of glyphosate in sediment at the pre-treatment timepoint in the Long Point pond (0.014 ppm) (Fig. 2D) – this concentration is below the limit of quantification and is therefore an estimate. In addition, there was a detection of glyphosate in the *P. australis* site at the pre-treatment timepoint in Turkey Point in 2017 (Fig. 2D). This was the result of an unlicensed, unaffiliated application of glyphosate to control *P. australis* (see Appendix G).

### Influence of Total Suspended Solids and Iron on Glyphosate and AMPA sediment concentrations

When controlling for distance from the point of application, total suspended solids (TSS) (mg/L) was not a strong predictor of the concentration of glyphosate in water (likelihood test ratio: X^2^ (1) = 0.179, p = 0.672), nor did it predict the concentration of AMPA in the water (likelihood test ratio: X^2^ (1) = 0.312, p = 0.577) (Appendix K). Similarly, total suspended solids (mg/L) and iron (mg/kg) did not predict the concentration of glyphosate in sediment (likelihood test ratio: X^2^ (2) = 1.065, p = 0.587) or the concentration of AMPA (likelihood test ratio: X^2^ (2) = 0.926, p = 0.629) (Appendix K) when distance from the point of application was controlled for. In both instances, the intercept model was a better fit and neither fixed factor improved the goodness of fit (Appendix L).

## Discussion

The risks associated with applying glyphosate-based herbicide directly over standing water to control invasive wetland plants are not well known (Breckels and Kilgour, 2018). Between 2016 and 2018, the Ontario government and partners began an unprecedented management program using a glyphosate-based herbicide to control invasive *Phragmites australis* was implemented in over 1000 ha of costal marsh on the North shore of Lake Erie. Despite extensive application, we did not observe glyphosate or AMPA concentrations in exceedance of concentrations of concern for aquatic biota in water or sediment at any timepoint. Dispersal of analytes typically remained within 10 m from the point of application, though trace amounts of glyphosate were collected more than 50 m from the site of application in both sediment and water. Glyphosate concentrations persisted longer in sediment than water samples, and evidence indicates concentrations return to pre-treatment levels at all stations after one-year.

### Glyphosate and AMPA

#### Maximum exposure risk

Concentrations of glyphosate and AMPA in water were highest within 24 hours of application but remained below the established CCME thresholds for the protection of aquatic life to short (27 ppm) or long-term (0.8 ppm) glyphosate exposure. Two application methods were used during the project, aerial and ground, with aerial application employing a higher loading rate of herbicide than ground application. The maximum concentrations of glyphosate in water detected after aerial application reflected this - there were more detections after aerial treatment which were on average higher than ground application, though the maximum concentrations were comparable. The concentrations of glyphosate in sediment followed the same pattern, with more detections that were on average higher within 24 hours of application, but similar maximum concentrations detected at the >20-day timepoint. These results indicate that one method does not pose a more significant risk to aquatic biota than the other.

Studies monitoring the concentration of glyphosate and AMPA in sediment are not as common as those that measure water. However, a survey of European agricultural topsoil from eleven countries, where glyphosate-based herbicide had been directly applied, found glyphosate and AMPA were present in just under half of their samples with a maximum glyphosate concentration of 2.00 ppm (AMPA approximately 1.50 ppm) (Silva et al., 2018). Battaglin et al. (2014) analysed 45 soil and sediment samples from Indiana and Mississippi and found a maximum concentration of 0.467 ppm for glyphosate and 0.341 ppm for AMPA. The concentrations of glyphosate and AMPA detected in the sediment from the marshes immediately after treatment were lower than these values, indicating direct application of glyphosate in a marsh does not result in the same level of sediment contamination as repeated applications in an agricultural field.

Concentrations of glyphosate we detected in water were often higher than those reported in environmental monitoring studies in North America, which we attribute to the direct over-water application that took place in our system. The maximum concentration we detected was 0.32 ppm, and our results were most similar to surface water in agriculturally dominated areas, such as the surface water assessment conducted by the USGS Toxic Substances Hydrology Program in nine midwestern states which detected a maximum glyphosate concentration of 0.417 ppm (0.041 ppm AMPA) (Scribner et al., 2007). In Southern Ontario streams, the majority of which were situated in watersheds dominated by agriculture or infrastructure, Struger et al. (2015) measured a maximum glyphosate concentration of 0.042 ppm (0.015 ppm for AMPA). Similar work monitoring surface waters in urban and rural watersheds in Ontario reported maximum glyphosate concentration of 0.012 ppm (Byer et al., 2008) and a Canada-wide assessment of urban rivers found a maximum glyphosate concentration of 0.012 ppm (Glozier et al., 2012). In rivers, streams, and wetlands in Southern Ontario the maximum glyphosate concentration detected was 0.041 ppm (0.066 ppm AMPA) (Struger et al., 2008) while lakes, ponds and wetlands in a U.S. study had a maximum glyphosate concentration of 0.301 ppm (0.041 ppm AMPA) (Battaglin et al., 2014). The results from these monitoring studies indicate that, while glyphosate and AMPA are ubiquitous in urban and rural watersheds, environmental concentrations are typically an order of magnitude below the maximum concentrations we observed following direct application to standing water. That said, our results are equivalent to some of the higher concentrations reported from environmental surface water samples collected from Canada and the US. Unfortunately, monitoring studies rarely provide sufficient covariate information to inform a direct comparison to published values.

### Dispersal and Persistence

Glyphosate and AMPA concentrations in water and sediment were highest closer to the point of herbicide application (i.e., within 10 m of treated *P. australis*), but trace concentrations of glyphosate were detected in water and sediment over 100 m from a point of application. AMPA was not detected outside areas of maximum exposure. In terms of persistence, glyphosate and AMPA in water generally returned to pre-treatment concentrations by the >20-day sampling timepoint. In contrast, glyphosate concentrations in sediment were higher at the >20-day timepoint and glyphosate was detectable in sediment up to one-year after treatment, though at lower concentrations. Once introduced into a system, glyphosate can be eliminated by one of two pathways: transport out of the area, or microbial decomposition (Borggaard and Gimsing, 2008).

Glyphosate has a high affinity for sediment (Borggaard and Gimsing, 2008; Grunewald et al., 2001) and adsorbs strongly to sandy soils (Aronsson et al., 2011). The sediment in Long Point, where the majority of our work took place, is comprised primarily of sand (≥ 88.4%) (Rooney lab, unpublished data). Once adsorbed, glyphosate persists in sediment, with a soil half-life that ranges from 2 to 215 days (Giesy et al., 2000). These properties of glyphosate explain why we detected it persisting in sediment in trace amounts for over a year. Sediment can then be re-suspended and transported out of the area, particularly in turbid aquatic systems, or it may settle to the bottom (e.g. Goldsborough and Brown, 1993). There, the adsorption and degradation of glyphosate depends on soil composition and properties (Borggaard and Gimsing, 2008). Though we did not detect a significant relationship between glyphosate concentration and iron in our system, there is a well-documented positive relationship between adsorption and iron oxides in sediment (Gimsing et al., 2004; Morillo et al., 2000; Ololade et al., 2014). The high affinity of glyphosate for sandy sediment likely explains the patterns of glyphosate dispersal and persistence that we saw, whereby glyphosate adsorbed rapidly from the water to suspended sediment and then settled to form a thin veneer of contaminated sediment. With wind and wave action, this veneer of sediment may be resuspended and deposited in new areas, leading to the spatial concentration of contaminated sediment in depositional areas.

The other pathway for eliminating glyphosate from a system is microbial decomposition. Microbes rapidly degrade glyphosate to AMPA (Sviridov et al., 2015), and the rate of microbial decomposition is inversely related to the sorption capacity of sediment (i.e. glyphosate becomes less bioavailable) (Borggaard and Gimsing, 2008). AMPA is water soluble and degrades more slowly than glyphosate (Grunewald et al., 2001), with a reported soil half-life that ranges from 60 – 240 days (Grandcoin et al., 2017). The low frequency of detection and generally low concentration of AMPA in our water and sediment samples reveal that decomposition of glyphosate by the AMPA pathway is relatively slow in our system, or that AMPA and glyphosate are rapidly flushed into the waters of the adjacent bays and diluted below detection limits.

### Aquasurf®

#### Maximum Exposure Risk, Dispersal and Persistence

Aquasurf® was rarely detected in water in 2016 and 2017 but was detected multiple times at high concentrations in 2018. Five of these samples exceeded the HERA alcohol ethoxylate (C_12_EO_6_) PNEC_water_ concentration of 0.129 ppm, three of which also exceeded the draft Canadian Federal Environmental Quality Guideline of 0.193 ppm. However, these detections were found at both the control and treatment transects and thus the origins of these alcohol ethoxylate homologues are unclear. Aquasurf® concentrations in sediment never approached thresholds of concern.

Aquasurf® was not detected often enough in the water and sediment among the three years to provide robust insight into its dispersal or persistence in the system. In 2018, there were more Aquasurf® detections than in previous years but they did not follow expected temporal or spatial pattern, including detections of Aquasurf® at stations along control transects where no herbicide was applied. For example, the high observation of Aquasurf® in water and sediment at the 50 m station was detected *prior* to herbicide. Further, the detected Aquasurf® was not associated with glyphosate or AMPA, and none of the intervening stations had any detectable levels of Aquasurf®. Alcohol ethoxylates, such as those in Aquasurf®, are anthropogenic in origin and could have arrived at these stations from other sources (e.g., soap residue washed off boats or discharged from cottages). Our monitoring cannot trace the detections to specific sources, but it does indicate that the herbicide-based control of *P. australis* was not the source. Despite no clear source of alcohol ethoxylates in surface water in 2018, the frequent detections indicate a potential area of concern for aquatic biota that warrants further study.

### Implications

Glyphosate-based herbicides, with an Aquasurf® surfactant, appears to be a safe and effective way to treat large scale invasions. Aerial application, when appropriate, minimizes trampling in sensitive ecosystems and while there are higher concentrations of glyphosate present right after treatment, the concentrations are still well below levels of concern. It is relevant, however, to acknowledge that the application of glyphosate can still lead to negative effects in aquatic ecosystems. With repeated applications, sediment can accumulate glyphosate and become a source of contamination (Battaglin et al., 2014; Myers et al., 2016) or it may accumulate in wetland vegetation (i.e. *Phragmites australis, Typha latifolia, Juncus effusus*) (Imfeld et al., 2013), where it can be released during decomposition. In addition, there are several reported sublethal effects of glyphosate, including delaying periphytic colonization, reducing the abundance of diatoms and encouraging the development of cyanobacteria (Vera et al., 2010). Low environmental levels can stimulate harmful algae growth (*Prymnesium parvum*) (Dabney and Patiño, 2018) and alter the timing of emergence of male and female tropical Chironomidae, which may have population level effects (Ferreira-Junior et al., 2017). These reports emphasize that the effects of low-level glyphosate exposure may be subtle and could result in consequences to aquatic ecosystems that we have not considered.

### Conclusions

- Direct over-water application of a custom, glyphosate-based herbicide posed no significant threat to aquatic biota in sensitive coastal marshes.
- While the loading rate of herbicide was higher with aerial application, the overall concentrations of glyphosate, AMPA, and Aquasurf® were comparable between the two methods.
- Glyphosate remains in the marsh sediment, at low concentrations, for over a year post-application.
- Alcohol ethoxylate detections did not follow expected patterns and cannot be attributed to Aquasurf® exposure. This highlights an area where further research is warranted.

## Funding

This work was supported by NSERC Discovery Grant #RGPIN-2014-03846 and MNRF non-consulting agreement MNRF-W-(12)3-16.

## Declaration of competing interests

There are no competing interests.

## Role of the funding source

Funding sources provided financial support for research technicians, graduate students, equipment and analysis. Funding sources did not have a role in study design, collection, analysis, or interpretation of data, in the writing of the report, or in the decision to submit the article for publication.

## Acknowledgements

We thank the technicians and graduate students who helped conduct field work: Graham Howell, Sarah Yuckin, Daina Anderson, Jessie Pearson, Heather Polowyk, Matthew Bolding, Jody Daniel, Jacob Basso, Laura Beecraft. A special thanks to *P. australis* control expert Dr. Janice Gilbert for her support. Thank you to the Ontario Ministry of Natural Resources and Forestry, the Ontario Ministry of Environment, Conservation and Parks, and the Canadian Wildlife Service for undertaking this extensive control work and supporting environmental monitoring. Thank you also to licensed herbicide applicator Eric Giles from Giles Restoration Services.

## Notes

### Competing Interest Statement

The authors have declared no competing interest.

