## Appendix for "Glyphosate used to control invasive *Phragmites australis* in standing water poses little risk to aquatic biota"

Appendix A. Dates of sampling in Rondeau Provincial Park, Long Point, and Turkey Point. Crown Marsh and Big Creek National Wildlife Area are managerial units within the Long Point region.

| Timepoint | Year | Marsh | Station | Date |
|---|---|---|---|---|
| Pre-treatment | 2016 | Rondeau PP | Ponds and transects | 1-Sept-2016 |
| | | Crown Marsh | | 2-Sept-2016 |
| 24 hours post treatment | 2016 | Rondeau PP | Ponds and transects | 15-Sept-2016 |
| | | Crown Marsh | | 16-Sept-2016 |
| >20 days post treatment | 2016 | Rondeau PP | Ponds and transects | 25-Oct-2016 |
| | | Crown Marsh | | 19-Oct-2016 |
| Pre-treatment | 2017 | Crown Marsh | | 15-Aug-2017 |
| | | Big Creek NWA | Transect | 17-Aug-2017 |
| | | Turkey Point | | 19-Aug-2017 |
| | | Crown Marsh | Re-sample 2016 sites | 20-Aug-2017 |
| | | Rondeau PP | | 25-Aug-2017 |
| 24 hours post treatment | 2017 | Crown Marsh | | 9-Sept-2017 |
| | | Big Creek NWA | Transect | 19-Sept-2017 |
| | | Turkey Point | | 13-Sept-2017 |
| >20 days post treatment | 2017 | Crown Marsh | | 26-Oct-2017 |
| | | Big Creek NWA | Transect | 22-Oct-2017 |
| | | Turkey Point | | 29-Oct-2017 |
| | | Crown Marsh | Re-sample 2016 sites | 7-Nov-2017 |
| Pre-treatment | 2018 | Crown Marsh | Follow-up transects | 1-Sept-2018 |
| | | Crown Marsh | Ponds, new treatment | 18-Sept-2018 |
| | | Turkey Point | Transect, new treatment | 1-Oct-2018 |
| | | Crown Marsh | Re-sample 2016 pond | 18-Sept-2018 |
| 24 hours post treatment | 2018 | Crown Marsh | Follow-up transects | 5-Sept-2018 |
| | | Crown Marsh | Ponds, new treatment | 23-Sept-2018 |
| | | Turkey Point | Transect, new treatment | 3-Oct-2018 |
| >20 days post treatment | 2018 | Crown Marsh | Follow-up transects | 9-Oct-2018 |
| | | Crown Marsh | Ponds, new treatment | 25-Oct-2018 |
| | | Turkey Point | Transect, new treatment | 25-Oct-2018 |

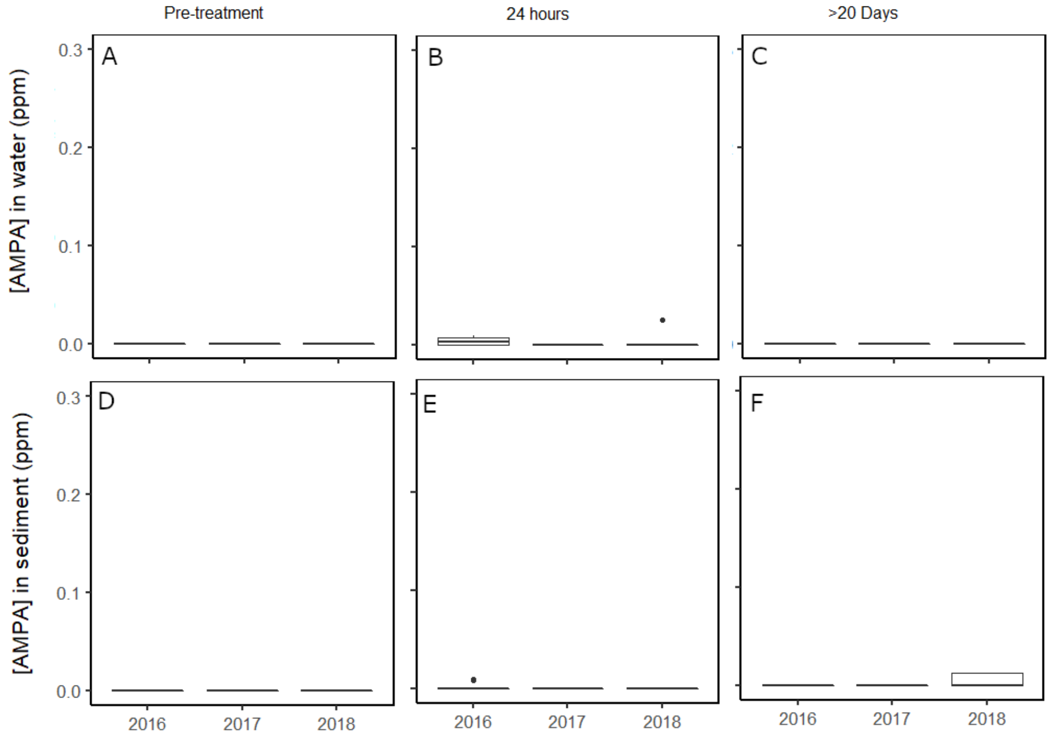


Appendix B. Concentrations of AMPA in water (A-C) and sediment (D-F) at sites with maximum exposure risk (i.e. *P. australis*, ponds, and 0 m transect sites) from treatment transects in Rondeau Provincial Park, Long Point, Big Creek National Wildlife Area, and Turkey Point, Ontario, CA (2016: n = 10, 2017: n = 6, 2018: n = 5). Boxplots represent the median, the 25th (lower hinge) and 95th (upper hinge) percentiles.

Appendix C. Average and maximum concentrations of glyphosate, AMPA and Aquasurf® detected in water samples at stations of maximum exposure (ponds, *Phragmites australis,* 0 m transect station). SE represents standard error. The detection limits for glyphosate and AMPA in water are 0.001 ppm and 0.002 ppm, respectively, and the limit of quantification for both is 0.008 ppm. For Aquasurf® the detection limit is 0.030 ppm and the limit of quantification is 0.080 ppm. Values below the limit of detection are reported as BDL here, and values below the limit of quantification represent certainty around presence but not the concentration of the analyte.

| Time | Year | Station | Glyphosate (ppm) | | | | AMPA (ppm) | | | | Aquasurf® surfactant (ppm) | | | |
|---|---|---|---|---|---|---|---|---|---|---|---|---|---|---|
| | | | Average | Max. | N | SE | Average | Max. | N | SE | Average | Max. | N | SE |
| Pre-treatment | 2016 | 0 m | BDL | BDL | 2 | BDL | BDL | BDL | 2 | BDL | BDL | BDL | 2 | BDL |
| | | *P. australis* | BDL | BDL | 2 | BDL | BDL | BDL | 2 | BDL | BDL | BDL | 2 | BDL |
| | | Pond | BDL | BDL | 6 | BDL | BDL | BDL | 6 | BDL | BDL | BDL | 6 | BDL |
| Pre-treatment | 2017 | 0 m | BDL | BDL | 3 | BDL | BDL | BDL | 3 | BDL | BDL | BDL | 3 | BDL |
| | | *P. australis* | BDL | BDL | 3 | BDL | BDL | BDL | 3 | BDL | BDL | BDL | 3 | BDL |
| Pre-treatment | 2018 | 0 m | BDL | BDL | 2 | BDL | BDL | BDL | 2 | BDL | BDL | BDL | 2 | BDL |
| | | *P. australis* | BDL | BDL | 2 | BDL | BDL | BDL | 2 | BDL | BDL | BDL | 2 | BDL |
| | | Pond | BDL | BDL | 1 | BDL | BDL | BDL | 1 | BDL | BDL | BDL | 1 | BDL |
| | | 0 m | 0.068 | 0.130 | 2 | 0.06 | 0.004 | 0.007 | 2 | 0.004 | BDL | BDL | 2 | BDL |
| 24 hours | 2016 | *P. australis* | 0.035 | 0.045 | 2 | 0.01 | 0.005 | 0.009 | 2 | 0.003 | BDL | BDL | 2 | BDL |
| | | Pond | 0.109 | 0.270 | 6 | 0.12 | 0.003 | 0.008 | 6 | 0.002 | 0.001 | 0.004 | 6 | 0.001 |
| 24 hours | 2017 | 0 m | 0.011 | 0.019 | 3 | 0.010 | BDL | BDL | 3 | BDL | BDL | BDL | 3 | BDL |
| | | *P. australis* | 0.008 | 0.011 | 3 | 0.002 | BDL | BDL | 3 | BDL | BDL | BDL | 3 | BDL |
| | | 0 m | 0.011 | 0.018 | 2 | 0.007 | BDL | BDL | 2 | BDL | BDL | BDL | 2 | BDL |

| Time | Year | Station | Glyphosate (ppm) | | | | AMPA (ppm) | | | | Aquasurf® surfactant (ppm) | | | |
|---|---|---|---|---|---|---|---|---|---|---|---|---|---|---|
| | | | Average | Max. | N | SE | Average | Max. | N | SE | Average | Max. | N | SE |
| 24 hours | 2018 | *P. australis* | 0.163 | 0.320 | 2 | 0.157 | 0.013 | 0.025 | 2 | 0.013 | 0.135 | 0.270 | 2 | 0.135 |
| | | Pond | 0.160 | 0.160 | 1 | - | BDL | BDL | 1 | - | BDL | BDL | 1 | - |
| | | 0 m | BDL | BDL | 2 | BDL | BDL | BDL | 2 | BDL | BDL | BDL | 2 | BDL |
| >20- days | 2016 | *P. australis* | 0.004 | 0.007 | 2 | 0.004 | BDL | BDL | 2 | BDL | 0.002 | 0.004 | 2 | 0.002 |
| | | Pond | BDL | BDL | 6 | BDL | BDL | BDL | 6 | BDL | BDL | BDL | 6 | BDL |
| >20-days | 2017 | 0 m | BDL | BDL | 3 | BDL | BDL | BDL | 3 | BDL | BDL | BDL | 3 | BDL |
| | | *P. australis* | BDL | BDL | 3 | BDL | BDL | BDL | 3 | BDL | BDL | BDL | 3 | BDL |
| >20-days | 2018 | 0 m | BDL | BDL | 2 | BDL | BDL | BDL | 2 | BDL | BDL | BDL | 2 | BDL |
| | | *P. australis* | BDL | BDL | 2 | BDL | BDL | BDL | 2 | BDL | BDL | BDL | 2 | BDL |
| | | Pond | BDL | BDL | 1 | - | BDL | BDL | 1 | - | BDL | BDL | 1 | BDL |

Appendix D. Average and maximum concentrations of glyphosate, AMPA and Aquasurf® detected in sediment samples at stations of maximum exposure. The calculated $PNEC_{soil}$ (predicted no effects concentration) for glyphosate (isopropylamine salt) in this system is 206.72 ppm. The detection limits for glyphosate and AMPA in sediment is 0.005 ppm, and the limit of quantification is 0.020 ppm. For Aquasurf® the detection limit is 0.300 ppm and the limit of quantification is 0.900 ppm. Values below the limit of detection are reported as BDL here, and values below the limit of quantification represent certainty around presence but not the concentration of the analyte.

| Time | Year | Station | Glyphosate (ppm) | | | | AMPA (ppm) | | | | Aquasurf® surfactant (ppm) | | | |
|---|---|---|---|---|---|---|---|---|---|---|---|---|---|---|
| | | | Average | Max. | N | SE | Average | Max. | N | SE | Average | Max. | N | SE |
| Pre-treatment | 2016 | 0 m | BDL | BDL | 2 | BDL | BDL | BDL | 2 | BDL | BDL | BDL | 2 | BDL |
| | | *P. australis* | BDL | BDL | 2 | BDL | BDL | BDL | 2 | BDL | BDL | BDL | 2 | BDL |
| | | Pond | 0.002 | 0.014 | 6 | 0.002 | BDL | BDL | 6 | BDL | BDL | BDL | 6 | BDL |
| Pre-treatment | 2017 | 0 m | BDL | BDL | 3 | BDL | BDL | BDL | 3 | BDL | BDL | BDL | 3 | BDL |
| | | *P. australis* | 0.005 | 0.016 | 3 | 0.005 | BDL | BDL | 3 | BDL | BDL | BDL | 3 | BDL |
| Pre-treatment | 2018 | 0 m | 0.002 | 0.005 | 2 | 0.002 | BDL | BDL | 2 | BDL | BDL | BDL | 2 | BDL |
| | | *P. australis* | 0.019 | 0.038 | 2 | 0.019 | BDL | BDL | 2 | BDL | BDL | BDL | 2 | BDL |
| | | Pond | BDL | BDL | 1 | - | BDL | BDL | 1 | BDL | BDL | BDL | 1 | BDL |
| 24 hours | 2016 | 0 m | 0.006 | 0.01 | 2 | 0.01 | BDL | BDL | 2 | BDL | BDL | BDL | 2 | BDL |

| Time | Year | Station | Glyphosate (ppm) | | | | AMPA (ppm) | | | | Aquasurf® surfactant (ppm) | | | |
|---|---|---|---|---|---|---|---|---|---|---|---|---|---|---|
| | | | Average | Max. | N | SE | Average | Max. | N | SE | Average | Max. | N | SE |
| | | *P. australis* | 0.060 | 0.12 | 2 | 0.06 | 0.004 | 0.009 | 2 | 0.004 | 0.7 | 1.4 | 2 | 0.7 |
| | | Pond | 0.036 | 0.11 | 6 | 0.02 | 0.002 | 0.01 | 6 | 0.002 | BDL | BDL | 6 | BDL |
| 24 hours | 2017 | 0 m | 0.009 | 0.02 | 3 | 0.005 | BDL | BDL | 3 | BDL | BDL | BDL | 3 | BDL |
| | | *P. australis* | 0.017 | 0.03 | 3 | 0.009 | BDL | BDL | 3 | BDL | BDL | BDL | 3 | BDL |
| 24 hours | 2018 | 0 m | 0.008 | 0.017 | 2 | 0.008 | BDL | BDL | 2 | BDL | BDL | BDL | 2 | BDL |
| | | *P. australis* | 0.012 | 0.024 | 2 | 0.012 | BDL | BDL | 2 | BDL | 0.6 | 1.2 | 2 | 0.6 |
| | | Pond | BDL | BDL | 1 | - | BDL | BDL | 1 | - | BDL | 0 | 1 | BDL |
| > 20-days | 2016 | 0 m | 0.014 | 0.03 | 2 | 0.014 | BDL | BDL | 2 | BDL | BDL | 0 | 2 | BDL |
| | | *P. australis* | 0.119 | 0.23 | 2 | 0.112 | BDL | BDL | 2 | BDL | 3.000 | 6.0 | 2 | 3.0 |
| | | Pond | 0.007 | 0.02 | 6 | 0.004 | BDL | BDL | 6 | BDL | BDL | BDL | 6 | BDL |
| > 20-days | 2017 | 0 m | BDL | BDL | 3 | BDL | BDL | BDL | 3 | BDL | BDL | BDL | 3 | BDL |
| | | *P. australis* | 0.019 | 0.056 | 3 | 0.019 | BDL | BDL | 3 | BDL | BDL | BDL | 3 | BDL |
| > 20-days | 2018 | 0 m | BDL | BDL | 2 | BDL | 0.006 | 0.013 | 2 | 0.006 | BDL | BDL | 2 | BDL |

| Time | Year | Station | Glyphosate (ppm) | | | | AMPA (ppm) | | | | Aquasurf® surfactant (ppm) | | | |
|---|---|---|---|---|---|---|---|---|---|---|---|---|---|---|
| | | | Average | Max. | N | SE | Average | Max. | N | SE | Average | Max. | N | SE |
| | | *P. australis* | 0.146 | 0.250 | 2 | 0.104 | 0.006 | 0.013 | 2 | 0.006 | BDL | BDL | 2 | BDL |
| | | Pond | BDL | BDL | 1 | BDL | BDL | BDL | 1 | - | BDL | BDL | 1 | BDL |

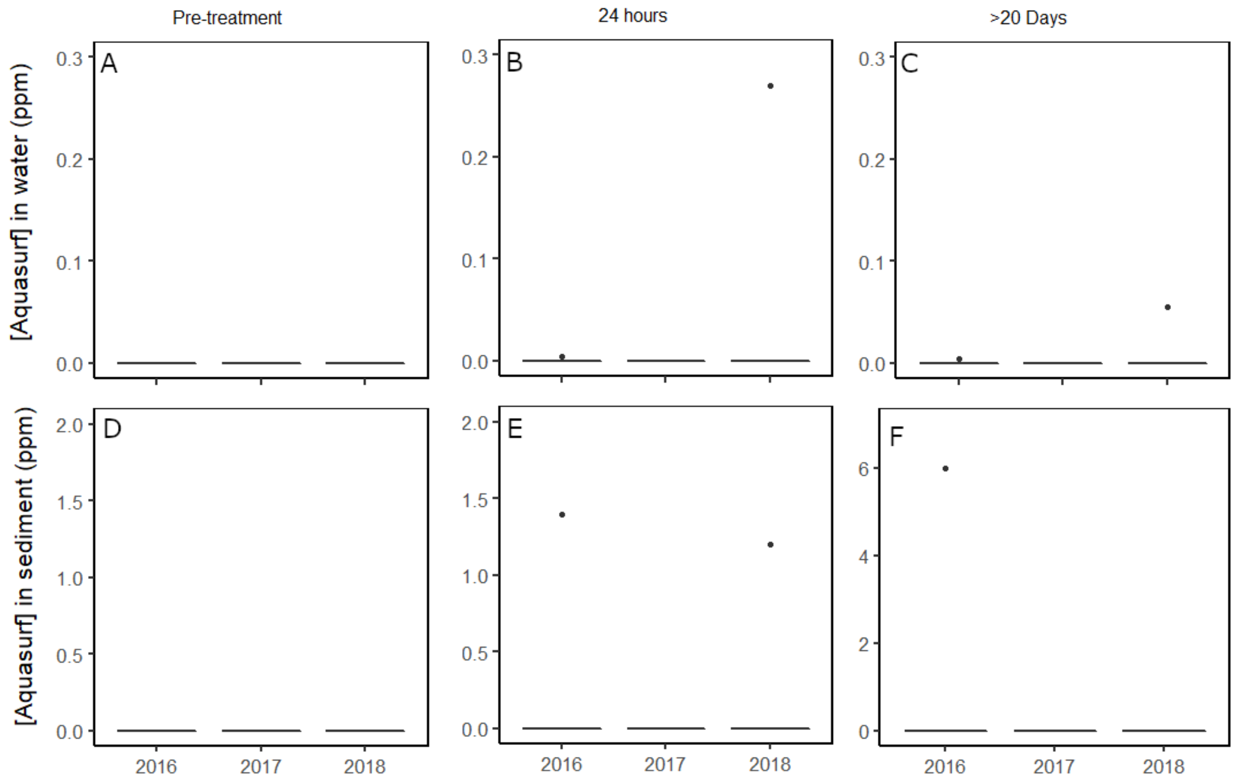


Appendix E. Concentrations of alcohol ethoxylates indicative of the surfactant Aquasurf® in water (A-C) and sediment (D-F) at sites with maximum exposure risk (i.e. *P. australis*, ponds, and 0 m transect sites) from treatment transects in Rondeau Provincial Park, Long Point, Big Creek National Wildlife Area, and Turkey Point, Ontario, CA (2016: n = 10, 2017: n = 6, 2018: n = 5). Note the difference in scale for Aquasurf® found in sediment >20 days later (F) which had a maximum of 6.00 ppm. Boxplots represent the median, the 25th (lower hinge) and 95th (upper hinge) percentiles.

Appendix F. Concentrations of glyphosate, its primary breakdown product aminomethylphosphonic acid (AMPA) and surfactant Aquasurf® in water samples. Aquasurf® here represents a collection of alcohol ethoxylate homologues that best represent the surfactant. Concentrations marked BDL are below the detection limit (0.001 ppm for glyphosate, 0.002 ppm for AMPA, and 0.030 ppm for Aquasurf®). Values in italics represent estimated concentrations, as they are above the detection limit but below the limit of quantification (0.008 ppm for glyphosate and AMPA) and Aquasurf® values below the limit of quantification (0.080 ppm) are represented as 0.600 ppm or halfway between the LOD and LOQ. Pond, *P. australis* (*Phragmites*) and 0 m transect station sites represent areas of maximum exposure. Samples were taken from stations in 2016, 2017, and 2018.

| Marsh | Year | Treatment | Timepoint | Station | Glyphosate (ppm) | AMPA (ppm) | Aquasurf® (ppm) |
|---|---|---|---|---|---|---|---|
| Long Point | 2016 | Herbicide | Pre-treatment | *Phragmites* | BDL | BDL | BDL |
| Long Point | 2016 | Herbicide | Pre-treatment | Pond | BDL | BDL | BDL |
| Long Point | 2016 | Herbicide | Pre-treatment | Pond | BDL | BDL | BDL |
| Long Point | 2016 | Herbicide | Pre-treatment | Pond | BDL | BDL | BDL |
| Long Point | 2016 | Herbicide | Pre-treatment | 0 m | BDL | BDL | BDL |
| Long Point | 2016 | Herbicide | Pre-treatment | 10 m | BDL | BDL | BDL |
| Long Point | 2016 | Herbicide | Pre-treatment | 50 m | BDL | BDL | BDL |
| Long Point | 2016 | Herbicide | Pre-treatment | 100 m | BDL | BDL | BDL |
| Long Point | 2016 | Herbicide | Pre-treatment | 150 m | BDL | BDL | BDL |
| Rondeau | 2016 | Herbicide | Pre-treatment | *Phragmites* | BDL | BDL | BDL |
| Rondeau | 2016 | Herbicide | Pre-treatment | Pond | BDL | BDL | BDL |
| Rondeau | 2016 | Herbicide | Pre-treatment | Pond | BDL | BDL | BDL |
| Rondeau | 2016 | Herbicide | Pre-treatment | Pond | BDL | BDL | BDL |
| Rondeau | 2016 | Herbicide | Pre-treatment | 0 m | BDL | BDL | BDL |
| Rondeau | 2016 | Herbicide | Pre-treatment | 10 m | BDL | BDL | BDL |
| Rondeau | 2016 | Herbicide | Pre-treatment | 50 m | BDL | BDL | BDL |
| Rondeau | 2016 | Herbicide | Pre-treatment | 100 m | BDL | BDL | BDL |
| Rondeau | 2016 | Herbicide | Pre-treatment | 150 m | BDL | BDL | BDL |
| Long Point | 2016 | Herbicide | 24 hours | *Phragmites* | 0.045 | 0.009 | BDL |
| Long Point | 2016 | Herbicide | 24 hours | Pond | 0.190 | *0.007* | *0.040* |
| Long Point | 2016 | Herbicide | 24 hours | Pond | 0.190 | *0.004* | BDL |
| Long Point | 2016 | Herbicide | 24 hours | Pond | 0.270 | 0.008 | BDL |
| Long Point | 2016 | Herbicide | 24 hours | 0 m | 0.130 | *0.007* | BDL |
| Long Point | 2016 | Herbicide | 24 hours | 10 m | 0.022 | BDL | BDL |
| Long Point | 2016 | Herbicide | 24 hours | 50 m | 0.011 | BDL | BDL |
| Long Point | 2016 | Herbicide | 24 hours | 100 m | BDL | BDL | BDL |
| Long Point | 2016 | Herbicide | 24 hours | 150 m | BDL | BDL | BDL |
| Rondeau | 2016 | Herbicide | 24 hours | *Phragmites* | 0.025 | *0.002* | BDL |
| Rondeau | 2016 | Herbicide | 24 hours | Pond | BDL | BDL | BDL |
| Rondeau | 2016 | Herbicide | 24 hours | Pond | BDL | BDL | BDL |
| Rondeau | 2016 | Herbicide | 24 hours | Pond | *0.007* | BDL | BDL |
| Rondeau | 2016 | Herbicide | 24 hours | 0 m | *0.005* | BDL | BDL |
| Rondeau | 2016 | Herbicide | 24 hours | 10 m | *0.004* | BDL | BDL |
| Rondeau | 2016 | Herbicide | 24 hours | 50 m | *0.001* | BDL | BDL |
| Rondeau | 2016 | Herbicide | 24 hours | 100 m | *0.001* | BDL | BDL |

| Marsh | Year | Treatment | Timepoint | Station | Glyphosate (ppm) | AMPA (ppm) | Aquasurf® (ppm) |
|---|---|---|---|---|---|---|---|
| Rondeau | 2016 | Herbicide | 24 hours | 150 m | BDL | BDL | BDL |
| Long Point | 2016 | Herbicide | >20 day | *Phragmites* | BDL | BDL | BDL |
| Long Point | 2016 | Herbicide | >20 day | Pond | BDL | BDL | BDL |
| Long Point | 2016 | Herbicide | >20 day | Pond | BDL | BDL | BDL |
| Long Point | 2016 | Herbicide | >20 day | Pond | BDL | BDL | BDL |
| Long Point | 2016 | Herbicide | >20 day | 0 m | BDL | BDL | BDL |
| Long Point | 2016 | Herbicide | >20 day | 10 m | BDL | BDL | BDL |
| Long Point | 2016 | Herbicide | >20 day | 50 m | BDL | BDL | BDL |
| Long Point | 2016 | Herbicide | >20 day | 100 m | BDL | BDL | BDL |
| Long Point | 2016 | Herbicide | >20 day | 150 m | BDL | BDL | BDL |
| Rondeau | 2016 | Herbicide | >20 day | *Phragmites* | *0.007* | BDL | *0.040* |
| Rondeau | 2016 | Herbicide | >20 day | Pond | BDL | BDL | BDL |
| Rondeau | 2016 | Herbicide | >20 day | Pond | BDL | BDL | BDL |
| Rondeau | 2016 | Herbicide | >20 day | Pond | BDL | BDL | BDL |
| Rondeau | 2016 | Herbicide | >20 day | 0 m | BDL | BDL | BDL |
| Rondeau | 2016 | Herbicide | >20 day | 10 m | BDL | BDL | BDL |
| Rondeau | 2016 | Herbicide | >20 day | 50 m | BDL | BDL | BDL |
| Rondeau | 2016 | Herbicide | >20 day | 100 m | BDL | BDL | BDL |
| Rondeau | 2016 | Herbicide | >20 day | 150 m | BDL | BDL | BDL |
| Big Creek | 2017 | Control | Pre-treatment | 100 m | BDL | BDL | BDL |
| Big Creek | 2017 | Control | Pre-treatment | 50 m | BDL | BDL | BDL |
| Big Creek | 2017 | Control | Pre-treatment | 25 m | BDL | BDL | BDL |
| Big Creek | 2017 | Control | Pre-treatment | 10 m | BDL | BDL | BDL |
| Big Creek | 2017 | Control | Pre-treatment | 0 m | BDL | BDL | BDL |
| Big Creek | 2017 | Control | Pre-treatment | *Phragmites* | BDL | BDL | BDL |
| Crown Marsh | 2017 | Control | Pre-treatment | 100 m | BDL | BDL | BDL |
| Crown Marsh | 2017 | Control | Pre-treatment | 50 m | BDL | BDL | BDL |
| Crown Marsh | 2017 | Control | Pre-treatment | 25 m | BDL | BDL | BDL |
| Crown Marsh | 2017 | Control | Pre-treatment | 10 m | BDL | BDL | BDL |
| Crown Marsh | 2017 | Control | Pre-treatment | 0 m | BDL | BDL | BDL |
| Crown Marsh | 2017 | Control | Pre-treatment | *Phragmites* | BDL | BDL | BDL |
| Turkey Point | 2017 | Control | Pre-treatment | 100 m | BDL | BDL | BDL |
| Turkey Point | 2017 | Control | Pre-treatment | 50 m | BDL | BDL | BDL |
| Turkey Point | 2017 | Control | Pre-treatment | 25 m | BDL | BDL | BDL |
| Turkey Point | 2017 | Control | Pre-treatment | 10 m | BDL | BDL | BDL |
| Turkey Point | 2017 | Control | Pre-treatment | 0 m | BDL | BDL | BDL |
| Turkey Point | 2017 | Control | Pre-treatment | *Phragmites* | BDL | BDL | BDL |
| Big Creek | 2017 | Control | 24 hours | 100 m | BDL | BDL | BDL |
| Big Creek | 2017 | Control | 24 hours | 50 m | BDL | BDL | BDL |
| Big Creek | 2017 | Control | 24 hours | 25 m | BDL | BDL | BDL |
| Big Creek | 2017 | Control | 24 hours | 10 m | BDL | BDL | BDL |
| Big Creek | 2017 | Control | 24 hours | 0 m | BDL | BDL | BDL |
| Big Creek | 2017 | Control | 24 hours | *Phragmites* | BDL | BDL | BDL |
| Crown Marsh | 2017 | Control | 24 hours | 100 m | BDL | BDL | BDL |
| Crown Marsh | 2017 | Control | 24 hours | 50 m | BDL | BDL | BDL |
| Crown Marsh | 2017 | Control | 24 hours | 25 m | BDL | BDL | BDL |
| Crown Marsh | 2017 | Control | 24 hours | 10 m | BDL | BDL | BDL |

| Marsh | Year | Treatment | Timepoint | Station | Glyphosate (ppm) | AMPA (ppm) | Aquasurf® (ppm) |
|---|---|---|---|---|---|---|---|
| Crown Marsh | 2017 | Control | 24 hours | 0 m | BDL | BDL | BDL |
| Crown Marsh | 2017 | Control | 24 hours | *Phragmites* | BDL | BDL | BDL |
| Turkey Point | 2017 | Control | 24 hours | 100 m | BDL | BDL | BDL |
| Turkey Point | 2017 | Control | 24 hours | 50 m | BDL | BDL | BDL |
| Turkey Point | 2017 | Control | 24 hours | 25 m | BDL | BDL | BDL |
| Turkey Point | 2017 | Control | 24 hours | 10 m | BDL | BDL | BDL |
| Turkey Point | 2017 | Control | 24 hours | 0 m | BDL | BDL | BDL |
| Turkey Point | 2017 | Control | 24 hours | *Phragmites* | BDL | BDL | BDL |
| Big Creek | 2017 | Control | >20 day | 100 m | BDL | BDL | BDL |
| Big Creek | 2017 | Control | >20 day | 50 m | BDL | BDL | BDL |
| Big Creek | 2017 | Control | >20 day | 25 m | BDL | BDL | BDL |
| Big Creek | 2017 | Control | >20 day | 10 m | BDL | BDL | BDL |
| Big Creek | 2017 | Control | >20 day | 0 m | BDL | BDL | BDL |
| Big Creek | 2017 | Control | >20 day | *Phragmites* | BDL | BDL | BDL |
| Crown Marsh | 2017 | Control | >20 day | 100 m | BDL | BDL | BDL |
| Crown Marsh | 2017 | Control | >20 day | 50 m | BDL | BDL | BDL |
| Crown Marsh | 2017 | Control | >20 day | 25 m | BDL | BDL | BDL |
| Crown Marsh | 2017 | Control | >20 day | 10 m | BDL | BDL | BDL |
| Crown Marsh | 2017 | Control | >20 day | 0 m | BDL | BDL | BDL |
| Crown Marsh | 2017 | Control | >20 day | *Phragmites* | BDL | BDL | BDL |
| Turkey Point | 2017 | Control | >20 day | 100 m | BDL | BDL | BDL |
| Turkey Point | 2017 | Control | >20 day | 50 m | BDL | BDL | BDL |
| Turkey Point | 2017 | Control | >20 day | 25 m | BDL | BDL | BDL |
| Turkey Point | 2017 | Control | >20 day | 10 m | BDL | BDL | BDL |
| Turkey Point | 2017 | Control | >20 day | 0 m | BDL | BDL | BDL |
| Turkey Point | 2017 | Control | >20 day | *Phragmites* | BDL | BDL | BDL |
| Big Creek | 2017 | Herbicide | Pre-treatment | 100 m | BDL | BDL | BDL |
| Big Creek | 2017 | Herbicide | Pre-treatment | 50 m | BDL | BDL | BDL |
| Big Creek | 2017 | Herbicide | Pre-treatment | 25 m | BDL | BDL | BDL |
| Big Creek | 2017 | Herbicide | Pre-treatment | 10 m | BDL | BDL | BDL |
| Big Creek | 2017 | Herbicide | Pre-treatment | 0 m | BDL | BDL | BDL |
| Big Creek | 2017 | Herbicide | Pre-treatment | *Phragmites* | BDL | BDL | BDL |
| Crown Marsh | 2017 | Herbicide | Pre-treatment | 100 m | BDL | BDL | BDL |
| Crown Marsh | 2017 | Herbicide | Pre-treatment | 50 m | BDL | BDL | BDL |
| Crown Marsh | 2017 | Herbicide | Pre-treatment | 25 m | BDL | BDL | BDL |
| Crown Marsh | 2017 | Herbicide | Pre-treatment | 10 m | BDL | BDL | BDL |
| Crown Marsh | 2017 | Herbicide | Pre-treatment | 0 m | BDL | BDL | BDL |
| Crown Marsh | 2017 | Herbicide | Pre-treatment | *Phragmites* | BDL | BDL | BDL |
| Turkey Point | 2017 | Herbicide | Pre-treatment | 100 m | BDL | BDL | BDL |
| Turkey Point | 2017 | Herbicide | Pre-treatment | 50 m | BDL | BDL | BDL |
| Turkey Point | 2017 | Herbicide | Pre-treatment | 25 m | BDL | BDL | BDL |
| Turkey Point | 2017 | Herbicide | Pre-treatment | 10 m | BDL | BDL | BDL |
| Turkey Point | 2017 | Herbicide | Pre-treatment | 0 m | BDL | BDL | BDL |
| Turkey Point | 2017 | Herbicide | Pre-treatment | *Phragmites* | BDL | BDL | BDL |
| Big Creek | 2017 | Herbicide | 24 hours | 100 m | BDL | BDL | BDL |
| Big Creek | 2017 | Herbicide | 24 hours | 50 m | BDL | BDL | BDL |
| Big Creek | 2017 | Herbicide | 24 hours | 25 m | BDL | BDL | BDL |

| Marsh | Year | Treatment | Timepoint | Station | Glyphosate (ppm) | AMPA (ppm) | Aquasurf® (ppm) |
|---|---|---|---|---|---|---|---|
| Big Creek | 2017 | Herbicide | 24 hours | 10 m | BDL | BDL | BDL |
| Big Creek | 2017 | Herbicide | 24 hours | 0 m | BDL | BDL | BDL |
| Big Creek | 2017 | Herbicide | 24 hours | *Phragmites* | *0.005* | BDL | BDL |
| Crown Marsh | 2017 | Herbicide | 24 hours | 100 m | BDL | BDL | BDL |
| Crown Marsh | 2017 | Herbicide | 24 hours | 50 m | *0.003* | BDL | BDL |
| Crown Marsh | 2017 | Herbicide | 24 hours | 25 m | 0.012 | BDL | BDL |
| Crown Marsh | 2017 | Herbicide | 24 hours | 10 m | *0.007* | BDL | BDL |
| Crown Marsh | 2017 | Herbicide | 24 hours | 0 m | 0.019 | BDL | BDL |
| Crown Marsh | 2017 | Herbicide | 24 hours | *Phragmites* | 0.011 | BDL | BDL |
| Turkey Point | 2017 | Herbicide | 24 hours | 100 m | BDL | BDL | BDL |
| Turkey Point | 2017 | Herbicide | 24 hours | 50 m | BDL | BDL | BDL |
| Turkey Point | 2017 | Herbicide | 24 hours | 25 m | *0.001* | BDL | BDL |
| Turkey Point | 2017 | Herbicide | 24 hours | 10 m | *0.008* | BDL | BDL |
| Turkey Point | 2017 | Herbicide | 24 hours | 0 m | 0.015 | BDL | BDL |
| Turkey Point | 2017 | Herbicide | 24 hours | *Phragmites* | *0.008* | BDL | BDL |
| Big Creek | 2017 | Herbicide | >20 day | 100 m | BDL | BDL | BDL |
| Big Creek | 2017 | Herbicide | >20 day | 50 m | BDL | BDL | BDL |
| Big Creek | 2017 | Herbicide | >20 day | 25 m | BDL | BDL | BDL |
| Big Creek | 2017 | Herbicide | >20 day | 10 m | BDL | BDL | BDL |
| Big Creek | 2017 | Herbicide | >20 day | 0 m | BDL | BDL | BDL |
| Big Creek | 2017 | Herbicide | >20 day | *Phragmites* | BDL | BDL | BDL |
| Crown Marsh | 2017 | Herbicide | >20 day | 100 m | BDL | BDL | *0.004* |
| Crown Marsh | 2017 | Herbicide | >20 day | 50 m | BDL | BDL | BDL |
| Crown Marsh | 2017 | Herbicide | >20 day | 25 m | BDL | BDL | BDL |
| Crown Marsh | 2017 | Herbicide | >20 day | 10 m | BDL | BDL | BDL |
| Crown Marsh | 2017 | Herbicide | >20 day | 0 m | BDL | BDL | BDL |
| Crown Marsh | 2017 | Herbicide | >20 day | *Phragmites* | BDL | BDL | BDL |
| Turkey Point | 2017 | Herbicide | >20 day | 100 m | BDL | BDL | BDL |
| Turkey Point | 2017 | Herbicide | >20 day | 50 m | BDL | BDL | BDL |
| Turkey Point | 2017 | Herbicide | >20 day | 25 m | BDL | BDL | BDL |
| Turkey Point | 2017 | Herbicide | >20 day | 10 m | BDL | BDL | BDL |
| Turkey Point | 2017 | Herbicide | >20 day | 0 m | BDL | BDL | BDL |
| Turkey Point | 2017 | Herbicide | >20 day | *Phragmites* | BDL | BDL | BDL |
| Crown Marsh | 2018 | Control | Pre-treatment | 100 m | BDL | BDL | BDL |
| Crown Marsh | 2018 | Control | Pre-treatment | 50 m | BDL | BDL | BDL |
| Crown Marsh | 2018 | Control | Pre-treatment | 25 m | BDL | BDL | BDL |
| Crown Marsh | 2018 | Control | Pre-treatment | 10 m | BDL | BDL | BDL |
| Crown Marsh | 2018 | Control | Pre-treatment | 0 m | BDL | BDL | BDL |
| Crown Marsh | 2018 | Control | Pre-treatment | *Phragmites* | BDL | BDL | BDL |
| Crown Marsh | 2018 | Herbicide | Pre-treatment | 100 m | BDL | BDL | BDL |
| Crown Marsh | 2018 | Herbicide | Pre-treatment | 50 m | BDL | BDL | BDL |
| Crown Marsh | 2018 | Herbicide | Pre-treatment | 25 m | BDL | BDL | BDL |
| Crown Marsh | 2018 | Herbicide | Pre-treatment | 10 m | BDL | BDL | BDL |
| Crown Marsh | 2018 | Herbicide | Pre-treatment | 0 m | BDL | BDL | BDL |
| Crown Marsh | 2018 | Herbicide | Pre-treatment | *Phragmites* | BDL | BDL | BDL |
| Crown Marsh | 2018 | Control | 24 hours | 100 m | BDL | BDL | BDL |
| Crown Marsh | 2018 | Control | 24 hours | 50 m | BDL | BDL | BDL |

| Marsh | Year | Treatment | Timepoint | Station | Glyphosate (ppm) | AMPA (ppm) | Aquasurf® (ppm) |
|---|---|---|---|---|---|---|---|
| Crown Marsh | 2018 | Control | 24 hours | 25 m | BDL | BDL | BDL |
| Crown Marsh | 2018 | Control | 24 hours | 10 m | BDL | BDL | BDL |
| Crown Marsh | 2018 | Control | 24 hours | 0 m | BDL | BDL | BDL |
| Crown Marsh | 2018 | Control | 24 hours | *Phragmites* | *0.001* | BDL | BDL |
| Crown Marsh | 2018 | Herbicide | 24 hours | 100 m | *0.002* | BDL | BDL |
| Crown Marsh | 2018 | Herbicide | 24 hours | 50 m | *0.002* | BDL | BDL |
| Crown Marsh | 2018 | Herbicide | 24 hours | 25 m | *0.002* | BDL | BDL |
| Crown Marsh | 2018 | Herbicide | 24 hours | 10 m | *0.002* | BDL | BDL |
| Crown Marsh | 2018 | Herbicide | 24 hours | 0 m | *0.004* | BDL | BDL |
| Crown Marsh | 2018 | Herbicide | 24 hours | *Phragmites* | *0.006* | BDL | BDL |
| Crown Marsh | 2018 | Herbicide | >20 day | 100 m | BDL | BDL | *0.055* |
| Crown Marsh | 2018 | Herbicide | >20 day | 50 m | BDL | BDL | *0.055* |
| Crown Marsh | 2018 | Herbicide | >20 day | 25 m | BDL | BDL | 0.130 |
| Crown Marsh | 2018 | Herbicide | >20 day | 10 m | BDL | BDL | *0.055* |
| Crown Marsh | 2018 | Herbicide | >20 day | 0 m | BDL | BDL | *0.055* |
| Crown Marsh | 2018 | Herbicide | >20 day | *Phragmites* | BDL | BDL | BDL |
| Crown Marsh | 2018 | Control | >20 day | 100 m | BDL | BDL | BDL |
| Crown Marsh | 2018 | Control | >20 day | 50 m | BDL | BDL | *0.055* |
| Crown Marsh | 2018 | Control | >20 day | 25 m | BDL | BDL | 0.120 |
| Crown Marsh | 2018 | Control | >20 day | 10 m | BDL | BDL | 0.110 |
| Crown Marsh | 2018 | Control | >20 day | 0 m | BDL | BDL | 2.500 |
| Crown Marsh | 2018 | Control | >20 day | *Phragmites* | BDL | BDL | BDL |
| Turkey Point | 2018 | Herbicide | Pre-treatment | 50 m | BDL | BDL | 0.140 |
| Turkey Point | 2018 | Herbicide | Pre-treatment | 25 m | BDL | BDL | BDL |
| Turkey Point | 2018 | Herbicide | Pre-treatment | 10 m | BDL | BDL | BDL |
| Turkey Point | 2018 | Herbicide | Pre-treatment | 0 m | BDL | BDL | BDL |
| Turkey Point | 2018 | Herbicide | Pre-treatment | *Phragmites* | BDL | BDL | BDL |
| Turkey Point | 2018 | Herbicide | 24 hours | 50 m | BDL | BDL | 0.200 |
| Turkey Point | 2018 | Herbicide | 24 hours | 25 m | BDL | BDL | BDL |
| Turkey Point | 2018 | Herbicide | 24 hours | 10 m | *0.005* | BDL | BDL |
| Turkey Point | 2018 | Herbicide | 24 hours | 0 m | 0.018 | BDL | BDL |
| Turkey Point | 2018 | Herbicide | 24 hours | *Phragmites* | 0.320 | 0.025 | 0.270 |
| Turkey Point | 2018 | Herbicide | >20 day | 50 m | BDL | BDL | BDL |
| Turkey Point | 2018 | Herbicide | >20 day | 25 m | BDL | BDL | BDL |
| Turkey Point | 2018 | Herbicide | >20 day | 10 m | BDL | BDL | BDL |
| Turkey Point | 2018 | Herbicide | >20 day | 0 m | BDL | BDL | BDL |
| Turkey Point | 2018 | Herbicide | >20 day | *Phragmites* | BDL | BDL | BDL |
| Crown Marsh | 2018 | Herbicide | Pre-treatment | Pond | BDL | BDL | BDL |
| Crown Marsh | 2018 | Control | Pre-treatment | Pond | BDL | BDL | BDL |
| Crown Marsh | 2018 | Herbicide | 24 hours | Pond | 0.016 | BDL | BDL |
| Crown Marsh | 2018 | Control | 24 hours | Pond | BDL | BDL | BDL |
| Crown Marsh | 2018 | Herbicide | >20 day | Pond | BDL | BDL | BDL |
| Crown Marsh | 2018 | Control | >20 day | Pond | BDL | BDL | BDL |

Appendix G. Concentrations of glyphosate, its primary breakdown product aminomethylphosphonic acid (AMPA) and surfactant Aquasurf® in sediment samples. Aquasurf® here represents a collection of alcohol ethoxylate homologues that best represent the surfactant. Concentrations marked BDL are below the detection limit (0.005 ppm for glyphosate and AMPA, and 0.300 ppm for Aquasurf®). Values in italics represent estimated concentrations, as they are above the detection limit but below the limit of quantification (0.020 ppm for glyphosate and AMPA) and Aquasurf® values below the limit of quantification (0.090 ppm) are represented as 0.600 ppm or halfway between the LOD and LOQ. Pond, *P. australis* (*Phragmites*) and 0 m transect station sites represent areas of maximum exposure. Samples were taken from stations in 2016, 2017, and 2018

| Marsh | Year | Treatment | Sample time | Distance | Glyphosate (ppm) | AMPA (ppm) | Aquasurf® (ppm) |
|---|---|---|---|---|---|---|---|
| Long Point | 2016 | Herbicide | Pre-treatment | *Phragmites* | BDL | BDL | BDL |
| Long Point | 2016 | Herbicide | Pre-treatment | Pond | *0.014* | BDL | BDL |
| Long Point | 2016 | Herbicide | Pre-treatment | Pond | BDL | BDL | BDL |
| Long Point | 2016 | Herbicide | Pre-treatment | Pond | BDL | BDL | BDL |
| Long Point | 2016 | Herbicide | Pre-treatment | 0 m | BDL | BDL | BDL |
| Long Point | 2016 | Herbicide | Pre-treatment | 10 m | BDL | BDL | BDL |
| Long Point | 2016 | Herbicide | Pre-treatment | 50 m | BDL | BDL | BDL |
| Long Point | 2016 | Herbicide | Pre-treatment | 100 m | BDL | BDL | BDL |
| Long Point | 2016 | Herbicide | Pre-treatment | 150 m | BDL | BDL | BDL |
| Rondeau | 2016 | Herbicide | Pre-treatment | *Phragmites* | BDL | BDL | BDL |
| Rondeau | 2016 | Herbicide | Pre-treatment | Pond | BDL | BDL | BDL |
| Rondeau | 2016 | Herbicide | Pre-treatment | Pond | BDL | BDL | BDL |
| Rondeau | 2016 | Herbicide | Pre-treatment | Pond | BDL | BDL | BDL |
| Rondeau | 2016 | Herbicide | Pre-treatment | 0 m | BDL | BDL | BDL |
| Rondeau | 2016 | Herbicide | Pre-treatment | 10 m | BDL | BDL | BDL |
| Rondeau | 2016 | Herbicide | Pre-treatment | 50 m | BDL | BDL | BDL |
| Rondeau | 2016 | Herbicide | Pre-treatment | 100 m | BDL | BDL | BDL |
| Rondeau | 2016 | Herbicide | Pre-treatment | 150 m | BDL | BDL | BDL |
| Long Point | 2016 | Herbicide | 24 hours | *Phragmites* | 0.120 | *0.009* | 1.400 |
| Long Point | 2016 | Herbicide | 24 hours | Pond | 0.110 | *0.01* | BDL |
| Long Point | 2016 | Herbicide | 24 hours | Pond | *0.014* | BDL | BDL |
| Long Point | 2016 | Herbicide | 24 hours | Pond | 0.028 | BDL | BDL |
| Long Point | 2016 | Herbicide | 24 hours | 0 m | *0.011* | BDL | BDL |
| Long Point | 2016 | Herbicide | 24 hours | 10 m | *0.013* | BDL | BDL |
| Long Point | 2016 | Herbicide | 24 hours | 50 m | BDL | BDL | BDL |
| Long Point | 2016 | Herbicide | 24 hours | 100 m | BDL | BDL | BDL |
| Long Point | 2016 | Herbicide | 24 hours | 150 m | BDL | BDL | BDL |
| Rondeau | 2016 | Herbicide | 24 hours | *Phragmites* | BDL | BDL | BDL |
| Rondeau | 2016 | Herbicide | 24 hours | Pond | BDL | BDL | BDL |
| Rondeau | 2016 | Herbicide | 24 hours | Pond | BDL | BDL | BDL |
| Rondeau | 2016 | Herbicide | 24 hours | Pond | 0.066 | BDL | BDL |
| Rondeau | 2016 | Herbicide | 24 hours | 0 m | BDL | BDL | BDL |
| Rondeau | 2016 | Herbicide | 24 hours | 10 m | BDL | BDL | BDL |
| Rondeau | 2016 | Herbicide | 24 hours | 50 m | BDL | BDL | BDL |
| Rondeau | 2016 | Herbicide | 24 hours | 100 m | BDL | BDL | BDL |

| Marsh | Year | Treatment | Sample time | Distance | Glyphosate (ppm) | AMPA (ppm) | Aquasurf® (ppm) |
|---|---|---|---|---|---|---|---|
| Rondeau | 2016 | Herbicide | 24 hours | 150 m | BDL | BDL | BDL |
| Long Point | 2016 | Herbicide | >20 day | *Phragmites* | *0.007* | BDL | BDL |
| Long Point | 2016 | Herbicide | >20 day | Pond | 0.022 | BDL | BDL |
| Long Point | 2016 | Herbicide | >20 day | Pond | BDL | BDL | BDL |
| Long Point | 2016 | Herbicide | >20 day | Pond | *0.017* | BDL | BDL |
| Long Point | 2016 | Herbicide | >20 day | 0 m | 0.027 | BDL | BDL |
| Long Point | 2016 | Herbicide | >20 day | 10 m | BDL | BDL | BDL |
| Long Point | 2016 | Herbicide | >20 day | 50 m | BDL | BDL | BDL |
| Long Point | 2016 | Herbicide | >20 day | 100 m | BDL | BDL | BDL |
| Long Point | 2016 | Herbicide | >20 day | 150 m | *0.009* | BDL | BDL |
| Rondeau | 2016 | Herbicide | >20 day | *Phragmites* | 0.230 | BDL | 6.000 |
| Rondeau | 2016 | Herbicide | >20 day | Pond | BDL | BDL | BDL |
| Rondeau | 2016 | Herbicide | >20 day | Pond | BDL | BDL | BDL |
| Rondeau | 2016 | Herbicide | >20 day | Pond | BDL | BDL | BDL |
| Rondeau | 2016 | Herbicide | >20 day | 0 m | BDL | BDL | BDL |
| Rondeau | 2016 | Herbicide | >20 day | 10 m | BDL | BDL | BDL |
| Rondeau | 2016 | Herbicide | >20 day | 50 m | BDL | BDL | BDL |
| Rondeau | 2016 | Herbicide | >20 day | 100 m | BDL | BDL | BDL |
| Rondeau | 2016 | Herbicide | >20 day | 150 m | BDL | BDL | BDL |
| Big Creek | 2017 | Control | Pre-treatment | 100 m | BDL | BDL | BDL |
| Big Creek | 2017 | Control | Pre-treatment | 50 m | BDL | BDL | BDL |
| Big Creek | 2017 | Control | Pre-treatment | 25 m | BDL | BDL | BDL |
| Big Creek | 2017 | Control | Pre-treatment | 10 m | BDL | BDL | BDL |
| Big Creek | 2017 | Control | Pre-treatment | 0 m | BDL | BDL | BDL |
| Big Creek | 2017 | Control | Pre-treatment | *Phragmites* | BDL | BDL | BDL |
| Crown Marsh | 2017 | Control | Pre-treatment | 100 m | BDL | BDL | BDL |
| Crown Marsh | 2017 | Control | Pre-treatment | 50 m | BDL | BDL | BDL |
| Crown Marsh | 2017 | Control | Pre-treatment | 25 m | BDL | BDL | BDL |
| Crown Marsh | 2017 | Control | Pre-treatment | 10 m | BDL | BDL | BDL |
| Crown Marsh | 2017 | Control | Pre-treatment | 0 m | BDL | BDL | BDL |
| Crown Marsh | 2017 | Control | Pre-treatment | *Phragmites* | BDL | BDL | BDL |
| Turkey Point | 2017 | Control | Pre-treatment | 100 m | BDL | BDL | BDL |
| Turkey Point | 2017 | Control | Pre-treatment | 50 m | BDL | BDL | BDL |
| Turkey Point | 2017 | Control | Pre-treatment | 25 m | BDL | BDL | BDL |
| Turkey Point | 2017 | Control | Pre-treatment | 10 m | BDL | BDL | BDL |
| Turkey Point | 2017 | Control | Pre-treatment | 0 m | BDL | BDL | BDL |
| Turkey Point | 2017 | Control | Pre-treatment | *Phragmites* | *0.016* | BDL | BDL |
| Big Creek | 2017 | Control | 24 hours | 100 m | BDL | BDL | BDL |
| Big Creek | 2017 | Control | 24 hours | 50 m | BDL | BDL | BDL |
| Big Creek | 2017 | Control | 24 hours | 25 m | BDL | BDL | BDL |
| Big Creek | 2017 | Control | 24 hours | 10 m | BDL | BDL | BDL |
| Big Creek | 2017 | Control | 24 hours | 0 m | BDL | BDL | BDL |
| Big Creek | 2017 | Control | 24 hours | *Phragmites* | BDL | BDL | BDL |
| Crown Marsh | 2017 | Control | 24 hours | 100 m | BDL | BDL | BDL |
| Crown Marsh | 2017 | Control | 24 hours | 50 m | BDL | BDL | BDL |
| Crown Marsh | 2017 | Control | 24 hours | 25 m | BDL | BDL | BDL |
| Crown Marsh | 2017 | Control | 24 hours | 10 m | BDL | BDL | BDL |

| Marsh | Year | Treatment | Sample time | Distance | Glyphosate (ppm) | AMPA (ppm) | Aquasurf® (ppm) |
|---|---|---|---|---|---|---|---|
| Crown Marsh | 2017 | Control | 24 hours | 0 m | BDL | BDL | BDL |
| Crown Marsh | 2017 | Control | 24 hours | *Phragmites* | BDL | BDL | BDL |
| Turkey Point | 2017 | Control | 24 hours | 100 m | BDL | BDL | BDL |
| Turkey Point | 2017 | Control | 24 hours | 50 m | BDL | BDL | BDL |
| Turkey Point | 2017 | Control | 24 hours | 25 m | BDL | BDL | BDL |
| Turkey Point | 2017 | Control | 24 hours | 10 m | BDL | BDL | BDL |
| Turkey Point | 2017 | Control | 24 hours | 0 m | BDL | BDL | BDL |
| Turkey Point | 2017 | Control | 24 hours | *Phragmites* | 0.020 | BDL | BDL |
| Big Creek | 2017 | Control | >20 day | 100 m | BDL | BDL | BDL |
| Big Creek | 2017 | Control | >20 day | 50 m | BDL | BDL | BDL |
| Big Creek | 2017 | Control | >20 day | 25 m | BDL | BDL | BDL |
| Big Creek | 2017 | Control | >20 day | 10 m | BDL | BDL | BDL |
| Big Creek | 2017 | Control | >20 day | 0 m | BDL | BDL | BDL |
| Big Creek | 2017 | Control | >20 day | *Phragmites* | BDL | BDL | BDL |
| Crown Marsh | 2017 | Control | >20 day | 100 m | BDL | BDL | BDL |
| Crown Marsh | 2017 | Control | >20 day | 50 m | BDL | BDL | BDL |
| Crown Marsh | 2017 | Control | >20 day | 25 m | BDL | BDL | BDL |
| Crown Marsh | 2017 | Control | >20 day | 10 m | BDL | BDL | BDL |
| Crown Marsh | 2017 | Control | >20 day | 0 m | BDL | BDL | BDL |
| Crown Marsh | 2017 | Control | >20 day | *Phragmites* | BDL | BDL | BDL |
| Turkey Point | 2017 | Control | >20 day | 100 m | BDL | BDL | BDL |
| Turkey Point | 2017 | Control | >20 day | 50 m | BDL | BDL | BDL |
| Turkey Point | 2017 | Control | >20 day | 25 m | BDL | BDL | BDL |
| Turkey Point | 2017 | Control | >20 day | 10 m | BDL | BDL | BDL |
| Turkey Point | 2017 | Control | >20 day | 0 m | BDL | BDL | BDL |
| Turkey Point | 2017 | Control | >20 day | *Phragmites* | 0.027 | BDL | BDL |
| Big Creek | 2017 | Herbicide | Pre-treatment | 100 m | BDL | BDL | BDL |
| Big Creek | 2017 | Herbicide | Pre-treatment | 50 m | BDL | BDL | BDL |
| Big Creek | 2017 | Herbicide | Pre-treatment | 25 m | BDL | BDL | BDL |
| Big Creek | 2017 | Herbicide | Pre-treatment | 10 m | BDL | BDL | BDL |
| Big Creek | 2017 | Herbicide | Pre-treatment | 0 m | BDL | BDL | BDL |
| Big Creek | 2017 | Herbicide | Pre-treatment | *Phragmites* | BDL | BDL | BDL |
| Crown Marsh | 2017 | Herbicide | Pre-treatment | 100 m | BDL | BDL | BDL |
| Crown Marsh | 2017 | Herbicide | Pre-treatment | 50 m | BDL | BDL | BDL |
| Crown Marsh | 2017 | Herbicide | Pre-treatment | 25 m | BDL | BDL | BDL |
| Crown Marsh | 2017 | Herbicide | Pre-treatment | 10 m | BDL | BDL | BDL |
| Crown Marsh | 2017 | Herbicide | Pre-treatment | 0 m | BDL | BDL | BDL |
| Crown Marsh | 2017 | Herbicide | Pre-treatment | *Phragmites* | BDL | BDL | 1.100 |
| Turkey Point | 2017 | Herbicide | Pre-treatment | 100 m | BDL | BDL | BDL |
| Turkey Point | 2017 | Herbicide | Pre-treatment | 50 m | BDL | BDL | BDL |
| Turkey Point | 2017 | Herbicide | Pre-treatment | 25 m | BDL | BDL | BDL |
| Turkey Point | 2017 | Herbicide | Pre-treatment | 10 m | BDL | BDL | BDL |
| Turkey Point | 2017 | Herbicide | Pre-treatment | 0 m | BDL | BDL | BDL |
| Turkey Point | 2017 | Herbicide | Pre-treatment | *Phragmites* | BDL | BDL | BDL |
| Big Creek | 2017 | Herbicide | 24 hours | 100 m | BDL | BDL | BDL |
| Big Creek | 2017 | Herbicide | 24 hours | 50 m | BDL | BDL | BDL |
| Big Creek | 2017 | Herbicide | 24 hours | 25 m | BDL | BDL | BDL |

| Marsh | Year | Treatment | Sample time | Distance | Glyphosate (ppm) | AMPA (ppm) | Aquasurf® (ppm) |
|---|---|---|---|---|---|---|---|
| Big Creek | 2017 | Herbicide | 24 hours | 10 m | BDL | BDL | BDL |
| Big Creek | 2017 | Herbicide | 24 hours | 0 m | BDL | BDL | BDL |
| Big Creek | 2017 | Herbicide | 24 hours | *Phragmites* | BDL | BDL | BDL |
| Crown Marsh | 2017 | Herbicide | 24 hours | 100 m | BDL | BDL | BDL |
| Crown Marsh | 2017 | Herbicide | 24 hours | 50 m | *0.005* | BDL | BDL |
| Crown Marsh | 2017 | Herbicide | 24 hours | 25 m | *0.015* | BDL | BDL |
| Crown Marsh | 2017 | Herbicide | 24 hours | 10 m | *0.013* | BDL | BDL |
| Crown Marsh | 2017 | Herbicide | 24 hours | 0 m | *0.011* | BDL | BDL |
| Crown Marsh | 2017 | Herbicide | 24 hours | *Phragmites* | *0.019* | BDL | BDL |
| Turkey Point | 2017 | Herbicide | 24 hours | 100 m | BDL | BDL | BDL |
| Turkey Point | 2017 | Herbicide | 24 hours | 50 m | BDL | BDL | BDL |
| Turkey Point | 2017 | Herbicide | 24 hours | 25 m | *0.006* | BDL | BDL |
| Turkey Point | 2017 | Herbicide | 24 hours | 10 m | 0.0435 | BDL | BDL |
| Turkey Point | 2017 | Herbicide | 24 hours | 0 m | *0.017* | BDL | BDL |
| Turkey Point | 2017 | Herbicide | 24 hours | *Phragmites* | 0.0324 | BDL | BDL |
| Big Creek | 2017 | Herbicide | >20 day | 100 m | BDL | BDL | BDL |
| Big Creek | 2017 | Herbicide | >20 day | 50 m | BDL | BDL | BDL |
| Big Creek | 2017 | Herbicide | >20 day | 25 m | *0.007* | BDL | BDL |
| Big Creek | 2017 | Herbicide | >20 day | 10 m | BDL | BDL | BDL |
| Big Creek | 2017 | Herbicide | >20 day | 0 m | BDL | BDL | BDL |
| Big Creek | 2017 | Herbicide | >20 day | *Phragmites* | BDL | BDL | BDL |
| Crown Marsh | 2017 | Herbicide | >20 day | 100 m | BDL | BDL | BDL |
| Crown Marsh | 2017 | Herbicide | >20 day | 50 m | BDL | BDL | BDL |
| Crown Marsh | 2017 | Herbicide | >20 day | 25 m | BDL | BDL | BDL |
| Crown Marsh | 2017 | Herbicide | >20 day | 10 m | BDL | BDL | BDL |
| Crown Marsh | 2017 | Herbicide | >20 day | 0 m | BDL | BDL | BDL |
| Crown Marsh | 2017 | Herbicide | >20 day | *Phragmites* | 0.056 | BDL | BDL |
| Turkey Point | 2017 | Herbicide | >20 day | 100 m | BDL | BDL | BDL |
| Turkey Point | 2017 | Herbicide | >20 day | 50 m | BDL | BDL | BDL |
| Turkey Point | 2017 | Herbicide | >20 day | 25 m | BDL | BDL | BDL |
| Turkey Point | 2017 | Herbicide | >20 day | 10 m | BDL | BDL | BDL |
| Turkey Point | 2017 | Herbicide | >20 day | 0 m | BDL | BDL | BDL |
| Turkey Point | 2017 | Herbicide | >20 day | *Phragmites* | BDL | BDL | BDL |
| Crown Marsh | 2018 | Control | Pre-treatment | 100 m | BDL | BDL | BDL |
| Crown Marsh | 2018 | Control | Pre-treatment | 50 m | BDL | BDL | BDL |
| Crown Marsh | 2018 | Control | Pre-treatment | 25 m | BDL | BDL | BDL |
| Crown Marsh | 2018 | Control | Pre-treatment | 10 m | BDL | BDL | BDL |
| Crown Marsh | 2018 | Control | Pre-treatment | 0 m | BDL | BDL | BDL |
| Crown Marsh | 2018 | Control | Pre-treatment | *Phragmites* | BDL | BDL | BDL |
| Crown Marsh | 2018 | Herbicide | Pre-treatment | 100 m | BDL | BDL | BDL |
| Crown Marsh | 2018 | Herbicide | Pre-treatment | 50 m | BDL | BDL | BDL |
| Crown Marsh | 2018 | Herbicide | Pre-treatment | 25 m | BDL | BDL | BDL |
| Crown Marsh | 2018 | Herbicide | Pre-treatment | 10 m | BDL | BDL | BDL |
| Crown Marsh | 2018 | Herbicide | Pre-treatment | 0 m | *0.005* | BDL | BDL |
| Crown Marsh | 2018 | Herbicide | Pre-treatment | *Phragmites* | 0.038 | BDL | BDL |
| Crown Marsh | 2018 | Control | 24 hours | 100 m | BDL | BDL | BDL |
| Crown Marsh | 2018 | Control | 24 hours | 50 m | BDL | BDL | BDL |

| Marsh | Year | Treatment | Sample time | Distance | Glyphosate (ppm) | AMPA (ppm) | Aquasurf® (ppm) |
|---|---|---|---|---|---|---|---|
| Crown Marsh | 2018 | Control | 24 hours | 25 m | BDL | BDL | BDL |
| Crown Marsh | 2018 | Control | 24 hours | 10 m | BDL | BDL | BDL |
| Crown Marsh | 2018 | Control | 24 hours | 0 m | BDL | BDL | BDL |
| Crown Marsh | 2018 | Control | 24 hours | *Phragmites* | BDL | BDL | 5.000 |
| Crown Marsh | 2018 | Herbicide | 24 hours | 100 m | BDL | BDL | BDL |
| Crown Marsh | 2018 | Herbicide | 24 hours | 50 m | BDL | BDL | BDL |
| Crown Marsh | 2018 | Herbicide | 24 hours | 25 m | BDL | BDL | BDL |
| Crown Marsh | 2018 | Herbicide | 24 hours | 10 m | BDL | BDL | BDL |
| Crown Marsh | 2018 | Herbicide | 24 hours | 0 m | *0.017* | BDL | BDL |
| Crown Marsh | 2018 | Herbicide | 24 hours | *Phragmites* | 0.024 | BDL | BDL |
| Crown Marsh | 2018 | Control | >20 day | 100 m | BDL | BDL | BDL |
| Crown Marsh | 2018 | Control | >20 day | 50 m | BDL | BDL | BDL |
| Crown Marsh | 2018 | Control | >20 day | 25 m | BDL | BDL | BDL |
| Crown Marsh | 2018 | Control | >20 day | 10 m | BDL | BDL | BDL |
| Crown Marsh | 2018 | Control | >20 day | 0 m | BDL | BDL | BDL |
| Crown Marsh | 2018 | Control | >20 day | *Phragmites* | BDL | BDL | BDL |
| Crown Marsh | 2018 | Herbicide | >20 day | 100 m | BDL | BDL | BDL |
| Crown Marsh | 2018 | Herbicide | >20 day | 50 m | BDL | BDL | BDL |
| Crown Marsh | 2018 | Herbicide | >20 day | 25 m | BDL | BDL | BDL |
| Crown Marsh | 2018 | Herbicide | >20 day | 10 m | BDL | BDL | BDL |
| Crown Marsh | 2018 | Herbicide | >20 day | 0 m | BDL | BDL | BDL |
| Crown Marsh | 2018 | Herbicide | >20 day | Phrag | 0.042 | BDL | BDL |
| Turkey Point | 2018 | Herbicide | Pre-treatment | 50 m | BDL | BDL | *0.600* |
| Turkey Point | 2018 | Herbicide | Pre-treatment | 25 m | BDL | BDL | BDL |
| Turkey Point | 2018 | Herbicide | Pre-treatment | 10 m | BDL | BDL | BDL |
| Turkey Point | 2018 | Herbicide | Pre-treatment | 0 m | BDL | BDL | BDL |
| Turkey Point | 2018 | Herbicide | Pre-treatment | *Phragmites* | BDL | BDL | BDL |
| Turkey Point | 2018 | Herbicide | 24 hours | 50 m | BDL | BDL | BDL |
| Turkey Point | 2018 | Herbicide | 24 hours | 25 m | BDL | BDL | BDL |
| Turkey Point | 2018 | Herbicide | 24 hours | 10 m | BDL | BDL | BDL |
| Turkey Point | 2018 | Herbicide | 24 hours | 0 m | BDL | BDL | BDL |
| Turkey Point | 2018 | Herbicide | 24 hours | *Phragmites* | BDL | BDL | 1.200 |
| Turkey Point | 2018 | Herbicide | 20day | 50 m | BDL | BDL | BDL |
| Turkey Point | 2018 | Herbicide | >20 day | 25 m | BDL | BDL | 2.400 |
| Turkey Point | 2018 | Herbicide | >20 day | 10 m | BDL | BDL | BDL |
| Turkey Point | 2018 | Herbicide | >20 day | 0 m | BDL | *0.013* | BDL |
| Turkey Point | 2018 | Herbicide | >20 day | *Phragmites* | 0.250 | *0.013* | BDL |
| Crown Marsh | 2018 | Herbicide | Pre-treatment | Pond 2 | BDL | BDL | BDL |
| Crown Marsh | 2018 | Control | Pre-treatment | Pond 1 | BDL | BDL | *0.600* |
| Crown Marsh | 2018 | Control | 24 hours | Pond 1 | BDL | BDL | *0.600* |
| Crown Marsh | 2018 | Herbicide | 24 hours | Pond 2 | BDL | BDL | BDL |
| Crown Marsh | 2018 | Herbicide | >20 day | Pond 2 | BDL | BDL | BDL |
| Crown Marsh | 2018 | Control | >20 day | Pond 1 | BDL | BDL | BDL |

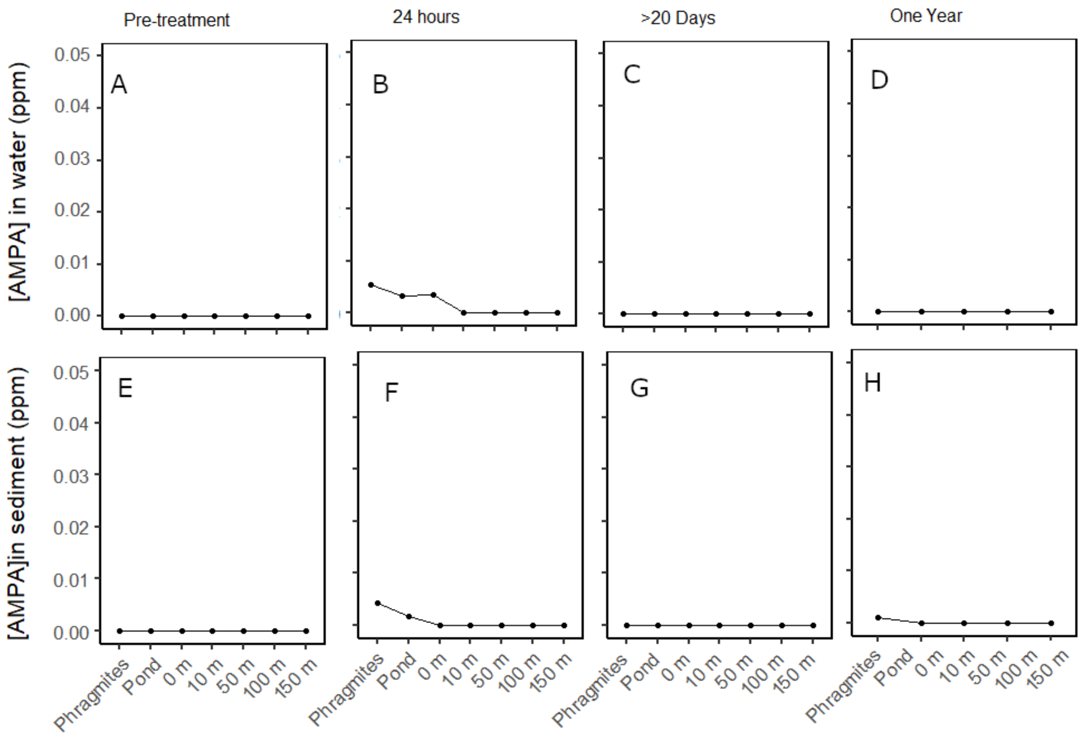


Appendix H. Concentrations of AMPA in water and sediment samples collected from Rondeau Provincial Park and Long Point, Ontario, CA. Samples were collected before aerial-application of glyphosate-based herbicide (pre-treatment), within 24 hours, >20-days after, and one-year after application. Sites represent areas of maximum exposure to herbicide (*P. australis*, ponds, 0 m) and distance (m) out into the adjoining bay. The average concentration at each point is represented here (pond: n = 6; Phragmites, 0, 10, 50, 100 and 150 m: n = 2). Data was collected in 2016 and 2017.

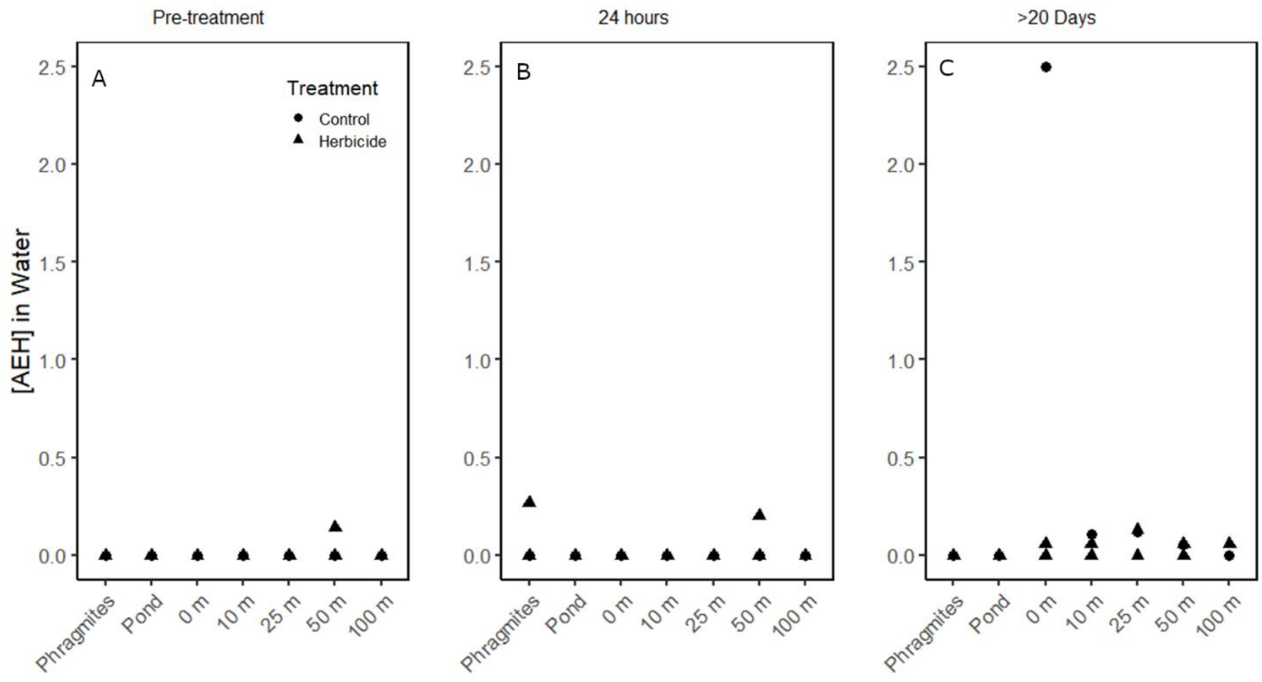


Appendix I. Detections of alcohol ethoxylates representative of the surfactant Aquasurf® in 2018. These samples did not follow an expected pattern of dispersal, as there were many detections in areas where herbicide was not applied (A, C). Samples were collected before treatment occurred (pre-treatment), within 24 hours of treatment, and >20 days after treatment from Crown Marsh and Turkey Point, Ontario, CA.

Appendix J. Concentrations of glyphosate, its primary breakdown product aminomethylphosphonic acid (AMPA) and surfactant Aquasurf® in water and sediment samples taken one to two years after initial glyphosate-based aerial application. Aquasurf® here represents a collection of alcohol ethoxylate homologues that best represent the surfactant. Concentrations marked BDL are below the detection limit.

| | | | | Water (ppm) | | Sediment (ppm) | |
|---|---|---|---|---|---|---|---|
| Marsh | Month | Timepoint | Station | Glyphosate | AMPA | Glyphosate | AMPA |
| Crown Marsh | Aug. 2017 | One year | *Phragmites* | BDL | BDL | 0.117 | 0.002 |
| Crown Marsh | Aug. 2017 | One year | Pond | BDL | BDL | BDL | BDL |
| Crown Marsh | Aug. 2017 | One year | 0 | BDL | BDL | BDL | BDL |
| Crown Marsh | Aug. 2017 | One year | 10 | BDL | BDL | BDL | BDL |
| Crown Marsh | Aug. 2017 | One year | 50 | BDL | BDL | BDL | BDL |
| Rondeau | Aug. 2017 | One year | *Phragmites* | BDL | BDL | BDL | BDL |
| Rondeau | Aug. 2017 | One year | Pond | BDL | BDL | BDL | BDL |
| Rondeau | Aug. 2017 | One year | 0 | BDL | BDL | BDL | BDL |
| Rondeau | Aug. 2017 | One year | 10 | BDL | BDL | BDL | BDL |
| Rondeau | Aug. 2017 | One year | 50 | BDL | BDL | BDL | BDL |
| Crown Marsh | Nov. 2017 | One year & 40 days | *Phragmites* | BDL | BDL | 0.094 | BDL |
| Crown Marsh | Nov. 2017 | One year & 40 days | Pond | BDL | BDL | BDL | BDL |
| Crown Marsh | Nov. 2017 | One year & 40 days | 0 | BDL | BDL | BDL | BDL |
| Crown Marsh | Nov. 2017 | One year & 40 days | 10 | BDL | BDL | BDL | BDL |
| Crown Marsh | Nov. 2017 | One year & 40 days | 50 | BDL | BDL | BDL | BDL |
| Rondeau | Nov. 2017 | One year & 40 days | *Phragmites* | BDL | BDL | BDL | BDL |
| Rondeau | Nov. 2017 | One year & 40 days | Pond | BDL | BDL | BDL | BDL |
| Rondeau | Nov. 2017 | One year & 40 days | 0 | BDL | BDL | BDL | BDL |
| Rondeau | Nov. 2017 | One year & 40 days | 10 | BDL | BDL | BDL | BDL |
| Rondeau | Nov. 2017 | One year & 40 days | 50 | BDL | BDL | BDL | BDL |
| Crown Marsh | Sept. 2018 | Two years | *Phragmites* | BDL | BDL | BDL | BDL |
| Crown Marsh | Sept. 2018 | Two years | Pond | BDL | BDL | BDL | BDL |
| Crown Marsh | Sept. 2018 | Two years | 0 | BDL | BDL | BDL | BDL |

Appendix K. Results of linear mixed effects models describing the relationship between glyphosate and AMPA concentrations (ppm) in water and total suspended solids (TSS) (mg/L), and glyphosate concentrations in sediment related to TSS and iron (Fe; mg/kg). Distance from the point of application was treated as a random effect in all models, and null models consisted of the intercept and random effect. Marginal ($r^2_m$) and conditional ($r^2_c$) coefficients of variation are reported to describe the proportion of variation explained by the fixed effects and the entire model, respectively. Error represents standard error.

| Model | Intercept ± error | Slope ± error | | t-value | | Goodness of Fit | |
|---|---|---|---|---|---|---|---|
| | | TSS | Fe | TSS | Fe | $r^2_m$ | $r^2_c$ |
| Glyphosate in water, null | 0.021 ± 0.012 | - | - | - | - | <0.001 | 0.199 |
| Glyphosate in sediment, null | <0.001 ± <0.001 | - | - | - | - | <0.001 | <0.001 |
| Glyphosate in water, full | 0.026 ± 0.016 | -0.007 ± 0.017 | - | -0.438 | - | 0.004 | 0.189 |
| Glyphosate in sediment, full | 0.002 ± 0.027 | 0.015 ± 0.015 | 0.005 ± 0.013 | 0.993 | 0.399 | 0.052 | 0.052 |
| AMPA in water, null | 0.001 ± <0.001 | - | - | - | - | <0.001 | 0.079 |
| AMPA in sediment, null | 0.001 ± 0.001 | - | - | - | - | <0.001 | <0.001 |
| AMPA in water, full | <0.001 ± 0.001 | <0.001 ± 0.001 | - | 0.561 | - | 0.007 | 0.090 |
| AMPA in sediment, full | <0.001 ± 0.001 | <0.001 ± 0.001 | <0.001 ± 0.001 | 0.950 | 0.281 | 0.045 | 0.045 |

Appendix L. AICc model comparisons were conducted for each pair of models (null and full) for each analyte (glyphosate, AMPA) and matrix (water, sediment) separately (pairs are shaded). In all cases, the null model was a better fit than the models that accounted for TSS (water) or TSS and Fe (sediment).

| Model | K | AICc | Δ | AICc weight |
|---|---|---|---|---|
| Glyphosate in water, null | 3 | -132.22 | 0.00 | 0.75 |
| Glyphosate in water, full | 4 | -130.00 | 2.22 | 0.25 |
| Glyphosate in sediment, null | 3 | -74.44 | 0.00 | 0.94 |
| Glyphosate in sediment, full | 5 | -68.94 | 5.52 | 0.06 |
| AMPA in water, null | 3 | -432.98 | 0.00 | 0.74 |
| AMPA in water, full | 4 | -421.89 | 2.09 | 0.26 |
| AMPA in sediment, null | 3 | -180.81 | 0.00 | 0.94 |
| AMPA in sediment, full | 5 | -175.15 | 5.66 | 0.06 |

Robichaud and Rooney. 2020. Glyphosate used to control invasive *Phragmites australis* in standing water poses little risk to aquatic biota. Submitted to Water Research 19 June 2020.
